# TCAB1 prevents nucleolar accumulation of the telomerase RNA to promote telomerase assembly

**DOI:** 10.1101/2021.05.27.445986

**Authors:** Basma M. Klump, Gloria I. Perez, Eric M. Patrick, Kate Adams-Boone, Scott B. Cohen, Li Han, Kefei Yu, Jens C. Schmidt

## Abstract

Localization of a wide variety of RNAs to non-membrane bound cellular compartments such as nucleoli, Cajal bodies, and stress-granules is critical for their function and stability. The molecular mechanisms that underly the recruitment and exclusion of specific RNAs from these phase-separated organelles is incompletely understood. Telomerase is a ribonucleoprotein (RNP) that is composed of the reverse transcriptase protein TERT, the telomerase RNA (TR), and several auxiliary proteins that associate with TR, including TCAB1. Here we show that in the absence of TCAB1, TR is tightly bound to the nucleolus, while TERT localizes to the nucleoplasm and is largely excluded from the nucleolus, significantly reducing telomerase assembly. Thus, nuclear compartmentalization by the non-membrane bound nucleolus counteracts telomerase assembly and TCAB1 is required to exclude the telomerase RNA from the nucleolus. Our work provides insight into the mechanism and functional consequences of RNA recruitment to organelles formed by phase separation proposes a new model explaining the critical role of TCAB1 in telomerase function.

## Introduction

Human cells contain a number of non-membrane bound organelles that carry out critical cellular functions. For instance, nucleoli and Cajal bodies are phase-separated nuclear organelles that play important roles in the biogenesis and maturation of many cellular RNAs (Hyman et al., 2014; Mitrea and Kriwacki, 2016). Nucleoli and Cajal bodies contain a wide range of small nucleolar and small Cajal body-specific RNAs (snoRNAs and scaRNAs, respectively). A subset of these snoRNAs and scaRNAs are bound by the H/ACA complex, which contains NOP10, NHP2, GAR1, and the pseudouridylase dyskerin, which modifies ribosomal and spliceosomal RNA precursors and other RNAs (Angrisani et al., 2014). A key difference between snoRNAs and scaRNAs is the presence of the Cajal-body box (CAB-box) motif in scaRNAs that directly associates with the telomerase Cajal body protein 1 (TCAB1, also known as WRAP53) (Jády et al., 2004; Schmidt and Cech, 2015; Venteicher et al., 2009). TCAB1 is required for the recruitment of scaRNAs to Cajal bodies and in its the absence scaRNAs localize to the nucleolus (Tycowski et al., 2009). Therefore, TCAB1 controls which phase-separated nuclear organelle scaRNAs associate with. Importantly, the molecular mechanism by which TCAB1 drives exclusion of scaRNAs from the nucleolus and facilitates their recruitment to Cajal bodies is unknown. In addition, it is unclear whether miss-localization of scaRNAs to the nucleolus has functional consequences.

The telomerase RNA (TR) is a scaRNA and, like other scaRNAs, its association with nucleoli and Cajal bodies is controlled by TCAB1 (Schmidt and Cech, 2015). Telomere maintenance by telomerase is essential for continuous proliferation of stem cell populations in the human body, and most cancers require telomerase activity for their survival (Stewart and Weinberg, 2006). To compensate for the incomplete replication of chromosome ends, telomerase appends TTAGGG repeats to the telomeric single-stranded overhang (Schmidt and Cech, 2015). Telomerase-mediated telomere maintenance requires three critical steps: telomerase assembly, telomerase recruitment to telomeres, and telomeric repeat synthesis (Schmidt and Cech, 2015). Mutations in several genes have been identified that cause deficiencies in one of these critical steps and lead to a variety of diseases known as telomere syndromes, which are characterized by premature depletion of stem cell populations (Armanios and Blackburn, 2012). In addition, telomerase is inappropriately activated in >85% of cancers (Stewart and Weinberg, 2006). While telomerase recruitment to telomeres (Nandakumar and Cech, 2013) and telomerase catalysis (Wu et al., 2017) have been studied extensively, much less is known about telomerase assembly. Importantly, telomerase assembly could be targeted to reduce telomerase activity in cancer cells, or to increase telomerase function in patients affected by genetically defined telomerase deficiency syndromes (Nagpal et al., 2020; Shukla et al., 2020).

Telomerase is a complex ribonucleoprotein (RNP). The core components of telomerase are the telomerase reverse transcriptase (TERT) protein, TR, the H/ACA complex, and TCAB1 (Schmidt and Cech, 2015). The primary function of the H/ACA complex is to stabilize TR, by directly binding to its 3’-end, preventing the exonucleolytic degradation of TR (Stuart et al., 2015; Tummala et al., 2015). The 3’-end formation of TR is tightly regulated by the competing activities of the poly-(A) polymerase PAPD5 and the nuclease PARN (Shukla et al., 2016; Tseng et al., 2015). Loss of TCAB1 function leads to telomere attrition in a variety of cell lines (Chen et al., 2018; Venteicher et al., 2009; Vogan et al., 2016; Zhong et al., 2011). In addition, multiple mutations in TCAB1 have been identified that cause misfolding of TCAB1 and lead to dyskeratosis congenita, a telomere syndrome (Freund et al., 2014; Zhong et al., 2011). While these observations highlight that TCAB1 is necessary for telomere synthesis, the underlying molecular mechanism is unclear. Initially, it was proposed that TCAB1 is required for telomerase recruitment to telomeres (Stern et al., 2012; Venteicher et al., 2009). A more recent study suggested that TCAB1 is required for the correct folding of TR, and that its absence causes a reduction in telomerase activity (Chen et al., 2018). All previous studies claim that TCAB1 does not mediate telomerase assembly, a critical step in telomere maintenance.

Here, we analyze telomerase assembly in intact cells and by purification of the telomerase RNP and demonstrate that, contrary to previous findings, TCAB1 promotes telomerase assembly *in vivo.* In the absence of TCAB1, TR is tightly associated with the nucleolus while TERT is largely excluded from the nucleolus. This spatial separation of TERT and TR, in the absence of TCAB1, is incompatible with proper telomerase assembly. Furthermore, we show that the limited amount of telomerase that can assemble in the absence of TCAB1 is fully active, suggesting that TCAB1 is not necessary for the enzymatic function of telomerase. Finally, analysis of the sub-cellular dynamics of TCAB1 revealed that it rarely enters the nucleolus, suggesting that TCAB1 associates with TR in the nucleoplasm and prevents its entry into the nucleolus rather than extracting TR molecules from the nucleolus. Collectively, our results support a model in which TCAB1 facilitates telomerase assembly by promoting nucleoplasmic accumulation of TR to increase encounters with TERT which is also localized to the nucleoplasm.

Furthermore, we demonstrate that the nucleolar phase separation constitutes a barrier for telomerase RNP assembly and suggest that incompletely assembled telomerase RNPs are tightly associated with the nucleolus and do not readily enter the nucleoplasm. Thus, cellular compartmentalization by phase-separated organelles such as the nucleolus can directly regulate RNP function in human cells.

## Results

### Loss of TCAB1 leads to nucleolar accumulation of TR

To assess whether TR is sequestered in the nucleolus in the absence of TCAB1, we knocked out TCAB1 in HeLa cells and HeLa cells expressing 3xFLAG-HaloTag-TERT (Halo-TERT) using Cas9 with two guide RNAs to delete exons 2 and 3 from the TCAB1 gene, which removes the coding sequence for residues 144-214 of TCAB1 and results in a frame shift (Fig. S1A). TCAB1 knock-out was validated by Southern blot, PCR, western blot, and immunofluorescence imaging (IF, Fig. 1A-C, Fig. S1B-C). To assure that no truncated form of TCAB1 was expressed in TCAB1 knock-out cells, we analyzed TCAB1 expression using two antibodies, targeting the N-terminus and C-terminus of TCAB1, respectively (Fig. S1D). Both HeLa and Halo-TERT TCAB1 knock-out cell lines continuously grew at approximately 60% of the rate of their parental cell lines (Fig. 1D). Telomeres in cells lacking TCAB1 were stable over time at a shorter length than telomeres in control cells, as previously described (Fig. S1E) (Vogan et al., 2016). Fluorescence *in situ* hybridization (FISH) demonstrated that TR accumulated in the nucleolus in cells that lack TCAB1, as indicated by co-localization of TR and nucleolar dyskerin signals (Fig. 1C). To confirm the dyskerin signal marks nucleoli, we transiently expressed HaloTag-dyskerin and GFP-NPM1, a well-established marker of the granular component (GC) of the nucleolus, in control and TCAB1 knock-out cells. Dyskerin was enriched in nucleolar regions with low NPM1 intensity in the presence and absence of TCAB1 (Fig. S1F), consistent with previously observed localization of dyskerin to the dense fibrillar component (DFC) of the nucleolus (Yao et al., 2019). Importantly, expression of GFP-TCAB1 in TCAB1 knock-out cells rescued TR localization to Cajal bodies (Fig. 1E), confirming that the miss-localization of TR to nucleoli is caused by absence of TCAB1. These observations demonstrate that TCAB1 is required to prevent TR accumulation in nucleoli.

**Figure 1.**
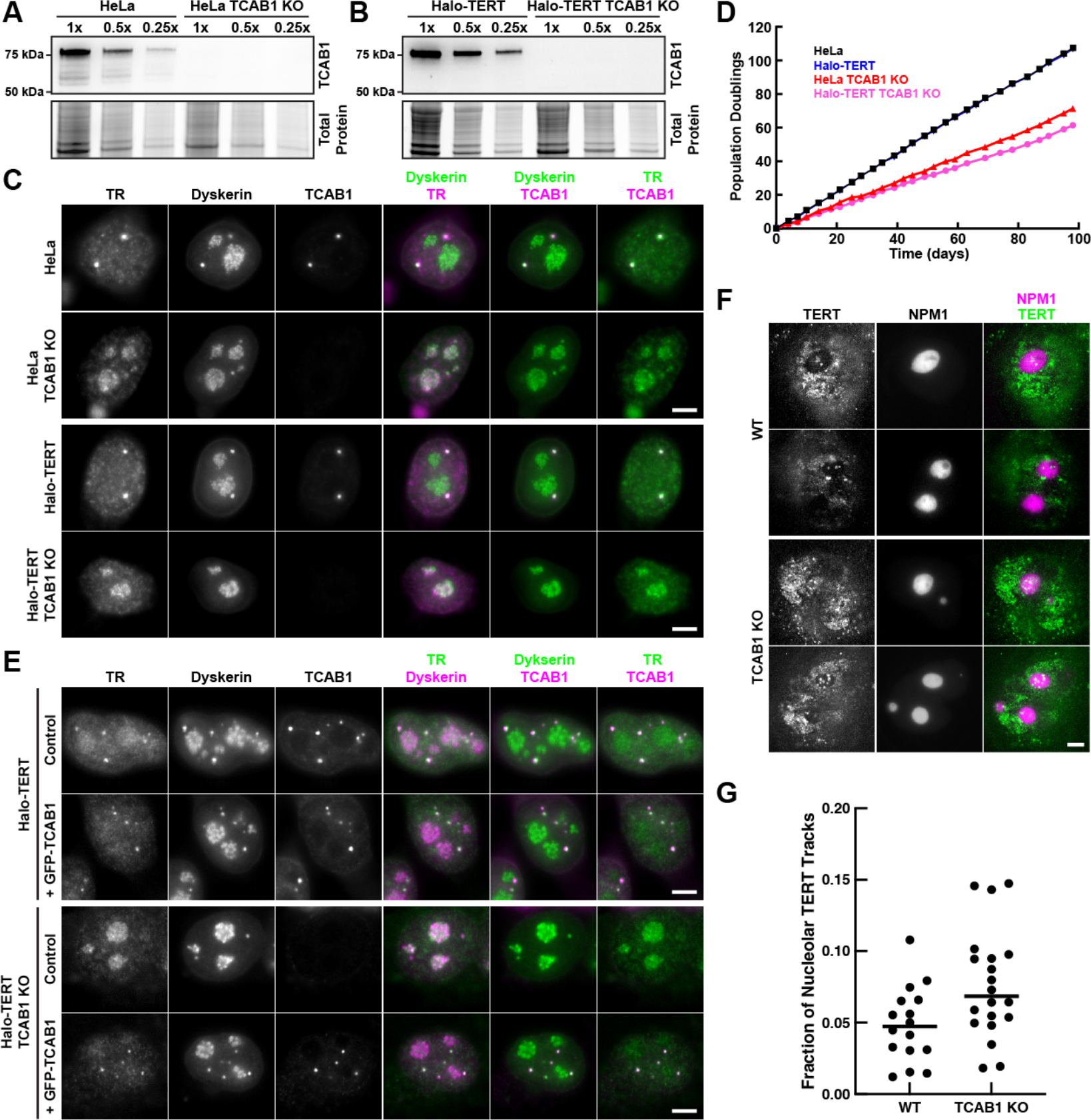
TR is localized to nucleoli in TCAB1 knock-out cells. **(A-B)** Western blot demonstrates the absence of TCAB1 protein in TCAB1 knock-out cell lines generated from **(A)** HeLa and **(B)** Halo-TERT parental cell lines (probed with Proteintech TCAB1 antibody). **(C)** Immuno-fluorescence with anti-dyskerin and anti-TCAB1 antibodies coupled to fluorescence in-situ hybridization with probes against TR, demonstrating the absence of TCAB1 and TR localization to nucleoli in TCAB1 knock-out cells (scale bar = 5 µm). **(D)** Growth rate of parental and TCAB1 knock-out cell lines. **(E)** Immuno-fluorescence with anti-dyskerin and anti-TCAB1 antibodies coupled to fluorescence in-situ hybridization with probes against TR, demonstrating that expression of GFP-TCAB1 in TCAB1 knock-out cells rescues TR localization to Cajal bodies (scale bar = 5 µm). **(F)** Maximum intensity projection (1000 frames at 10 ms per frame) of HaloTag-TERT (JFX650) in control and TCAB1 knock-out cells expressing GFP-NPM1 to mark nucleoli (scale bar = 5 µm). **(G)** Quantification of the fraction of TERT trajectories that overlap with the nucleolus in control and TCAB1 knock-out cells.

### TERT is not enriched in nucleoli in the absence of TCAB1

Our previous observations demonstrated that TERT does not enter nucleoli in wild type human cancer cells (Schmidt et al., 2016). To test whether TERT, like TR, is enriched in nucleoli in the absence of TCAB1, we performed single-molecule imaging of 3xFLAG-HaloTag-TERT in living HeLa and TCAB1 knock-out cell lines transiently expressing GFP-NPM1 as a nucleolar marker. 3xFLAG-HaloTag-TERT was largely excluded from the nucleolus in the presence and absence of TCAB1 with a small number of TERT molecules localizing to nucleoli marker by NPM1 (Fig. 1F, Movies S1-2). Single-particle tracking revealed that 4.9% and 7.7% of TERT trajectories overlapped with the nucleolus in control and TCAB1 knock-out cells, respectively (Fig. 1G). To exclude the possibility that nucleolar exclusion is a consequence of the 3xFLAG-HaloTag on the N-terminus of TERT used in our experiments, we transiently expressed the 3xFLAG-HaloTag fused to a nuclear localization sequence (NLS) in HeLa cells. Single-molecule imaging demonstrated that the nuclear 3xFLAG-HaloTag signals overlapped with the nucleolus (Fig. S1G-H, Movie S3). Similar to the 3xFLAG-HaloTag alone, 3xFLAG-HaloTag-dyskerin also localized to the nucleolus (Fig. S1F). These results demonstrate that 3xFLAG-HaloTag-TERT is largely excluded from the nucleolus in the presence and absence of TCAB1 and that this exclusion is not caused by the 3xFLAG-HaloTag but instead is an intrinsic property of TERT. In contrast, TR accumulates in the nucleolus in the absence of TCAB1 which demonstrates that the vast majority of TERT does not co-localize with TR in cells lacking TCAB1 and therefore is not associated with TR.

### TCAB1 promotes telomerase RNP assembly

Previous studies by other laboratories have concluded that telomerase assembly is unaffected by the absence of TCAB1 and whether TCAB1 is required for telomerase activity is controversial (Chen et al., 2018; Venteicher et al., 2009; Vogan et al., 2016). Importantly, telomerase assembly was only qualitatively assessed in these studies. To quantitatively analyze the role of TCAB1 in telomerase assembly, we immuno-purified endogenous telomerase using a well-established anti-TERT antibody (Cohen et al., 2007). The amount of TERT purified from TCAB1 knock-out cells was reduced compared to control cells (Fig. 2A), which likely indicates a lower expression level of TERT in cells lacking TCAB1. In contrast, TR levels were not reduced in cells lacking TCAB1 (Fig. 2B). To quantify telomerase assembly, we measured TERT levels by western blot and determined the amount of TR co-purified using Northern blot (Fig. 2A,B). The amount of TR associated with TERT relative to total cellular TR was reduced to <20% in TCAB1 knock-out cells compared to parental controls (Fig. 2B-C). In addition, the ratio of TR relative to TERT, which is a direct measure of telomerase assembly, was reduced to 20-40% in cells lacking TCAB1 relative to controls (Fig. 2A-B,D). This excludes the possibility that the lower amount of TR co-purified with TERT from TCAB1 knock-out cells is a consequence of the reduction of total TERT immuno-precipitated from these cells (Fig. 2A,D). These observations strongly suggest that telomerase assembly is defective in cells that lack TCAB1. To further test this hypothesis, we overexpressed TERT and TR in parental and TCAB1 knock-out cells and immuno-purified TERT with the same TERT antibody (Fig. 2E, Fig. S2A). After overexpression of TERT and TR, the fraction of TR associated with TERT and the ratio of TR relative to TERT were significantly reduced when telomerase was purified from TCAB1 knock-out cells (Fig. 2E-F,H-I, Fig. S2B,D). Furthermore, the increased amount of telomerase purified after over expression allowed us to assess the amount of dyskerin associated with the telomerase RNP (Fig. 2G, Fig. S2C). Since TR bridges TERT and dyskerin, dyskerin co-purified with TERT directly reports on the presence of TR. Consistent with the reduction in TR, the amount of dyskerin bound to TERT was also reduced to about 25% when TERT was purified from cells lacking TCAB1 compared to parental controls (Fig. 2G, H, Fig. S2C,E). Importantly, we also confirmed that TCAB1 was absent from telomerase purified from TCAB1 knock-out cells (Fig. 2E, Fig. S2A). Altogether, these results demonstrate that telomerase assembly is significantly reduced (to ∼20-40% of control levels) in the absence of TCAB1 and that overexpression of TERT and TR is not sufficient to overcome this defect in telomerase RNP formation.

**Figure 2.**
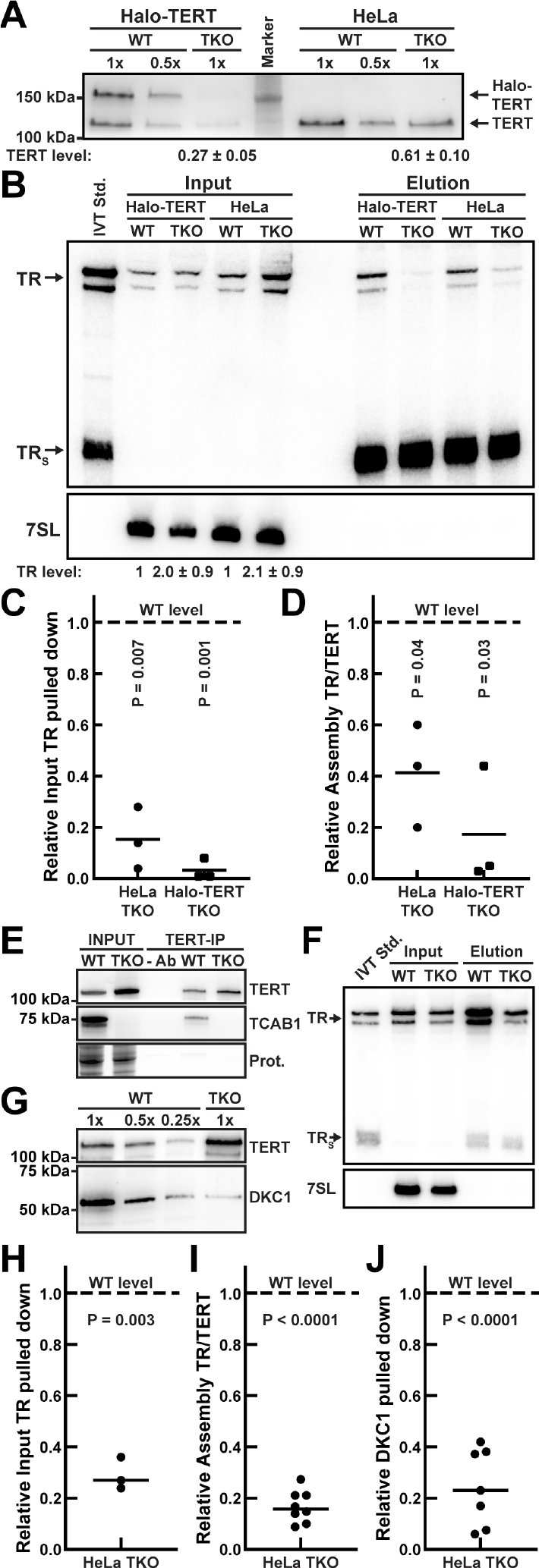
Telomerase Assembly is reduced in the absence of TCAB1. **(A)** Western blots analyzing endogenous TERT immuno-purification (using a sheep anti-TERT antibody) probed with a rabbit anti-TERT antibody (Abcam). TERT level normalized to WT (n = 3, SD). **(B)** Northern blot of RNA extracted from input and purified endogenous TERT samples probed with radiolabeled DNA oligonucleotides complementary to TR. Standards are *in vitro* transcribed full-length TR and truncated TRS. TRS was added to samples prior to RNA extraction as loading and recovery control. Input samples were also probed for 7SL RNA as loading control. Input TR levels relative to WT control normalized to 7SL RNA (n = 3, SD). **(C-D)** Quantification of the amount of **(C)** TR purified relative to input RNA levels, and **(D)** the ratio of TR relative to endogenous TERT in telomerase purified from TCAB1 knock-out cells compared to controls (n = 3, mean, T-Test). The dashed lines indicate the level in telomerase purified from wild-type TCAB1 cells which was normalized to 1.0. **(E-J)** TERT immuno-purification (using a sheep anti-TERT antibody) from HeLa control cells (WT) or TCAB1 knock-out cells (TKO) overexpressing TERT and TR. **(E)** Western blots of immuno-purified TERT probed with a rabbit anti-TERT antibody (Abcam) and an anti-TCAB1 antibody. **(F)** Northern blot of RNA extracted from input and purified TERT (Elution) probed with radiolabeled DNA oligonucleotides complementary to TR. Standards are *in vitro* transcribed full-length TR and truncated TRS. TRS was added to samples prior to RNA extraction as loading and recovery control. Input samples were also probed for 7SL RNA as loading control. **(G)** Western blots to analyze immuno-purified telomerase RNP composition. A single membrane was cut into two pieces that were probed with anti-TERT and anti-dyskerin antibodies, respectively. **(H-J)** Quantification of the amount of **(H)** TR relative to input TR (n = 3), **(I)** the ratio of TR to TERT (n = 8), and **(J)** dyskerin (n = 7) in TERT purifications from TCAB1 knock-out cells overexpressing TERT and TR compared to parental controls (mean, T-Test). The dashed lines indicate the level in telomerase purified from wild-type TCAB1 cells which was normalized to 1.0.

To assure that the reduction in telomerase assembly observed in TCAB1 knock-out cells is not a consequence of an altered cell cycle distribution of cells that lacked TCAB1, we purified telomerase from cells sychronized in various stages of the cell cycle. Due to the slow growth of the TCAB1 knock-out cells a double thymidine block was not feasible. We therefore synchronized cells using thymidine for 24 hours, released and harvested cells immediately (early S phase), 4 hours (mid S phase), and 8 hours (G2/M phase) after release. Cell-cycle synchronization was confirmed by DNA content analysis using propidium iodide staining and flow cytometry (Fig. S2F). Telomerase assembly was reduced in TCAB1 knock-out cells in all cell cycle stages analyzed (Fig. S2G-I). This demonstrates that the reduction of telomerase assembly observed in cells lacking TCAB1 is not a consequence of an aberant cell-cycle distribution of TCAB1 knock-out cells.

### TCAB1 is not required for telomerase catalytic activity

To assess whether TCAB1 plays a role in telomerase catalysis, we first analyzed the enzymatic activity of endogenous telomerase purified from TCAB1 knock-out cells using the direct telomerase extension assays (Fig. 3A,B). Consistent with previous results (Chen et al., 2018), telomerase activity was strongly reduced in the absence of TCAB1 (Fig. 3C). To address whether this reduction in telomerase activity was a consequence of the defect in telomerase assembly observed in TCAB1 knock-out cells, we determined the specific activity of telomerase by dividing the measured activity by the amount of TR present in the respective telomerase sample. Due to the very small amount of TR detected in endogenous telomerase samples (Fig. 2B-C), the specific activity of endogenous telomerase was highly variable in TCAB1 knock-out cells but did not appear to be reduced compared to telomerase purified from control cells (Fig. S4A). To overcome this limitation, we determined the specific activity of telomerase purified from cells overexpressing TERT and TR. Similar to endogenous telomerase, activity of overexpressed telomerase purified from HeLa and Halo-TERT cells lacking TCAB1 was significantly reduced to 24% and 34% compared to controls, respectively (Fig. 3D-F, Fig. S3). The specific activity of overexpressed telomerase purified from HeLa and Halo-TERT cells lacking TCAB1 was slightly reduced (84% and 77% relative to control, respectively), but this reduction was not statistically significant (Fig. 3G). Together these observations demonstrate that net cellular telomerase activity is reduced in the absence of TCAB1, while specific enzymatic activity is not. The reduction in cellular telomerase activity corresponds closely to the reduction of telomerase assembly observed in TCAB1 knock-out cells, suggesting that the smaller number of telomerase RNPs that form in the absence of TCAB1 are fully active.

**Figure 3.**
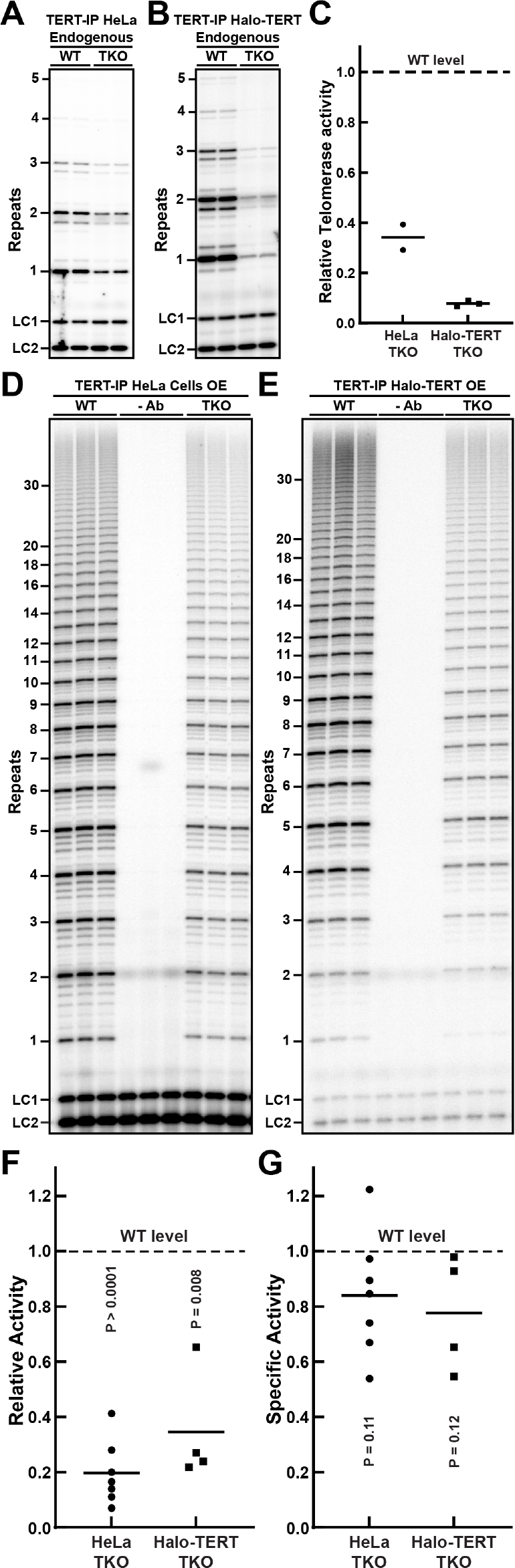
The specific activity of telomerase is unchanged in the absence of TCAB1. **(A-B)** Direct telomerase extension assay of endogenous telomerase immuno-purified from parental (WT) and TCAB1 knock-out (TKO) **(A)** HeLa and **(B)** Halo-TERT cell lines. Assays were carried out in the presence of 300 mM KCl and limiting amounts of dGTP. LC1 and LC2, radiolabeled DNA oligonucleotide loading controls. **(C)** Quantification of endogenous telomerase activity in samples from TCAB1 knock-out cells relative to parental controls (n = 3, mean). **(D-E)** Direct telomerase extension assay of overexpressed telomerase immuno-purified from parental (WT) and TCAB1 knock-out (TKO) **(D)** HeLa and **(E)** Halo-TERT cell lines. Assays were carried out in the presence of 150 mM KCl and 10 µM dATP, dTTP, and dGTP. LC1 and LC2, radiolabeled DNA oligonucleotide loading controls. In -Ab samples the TERT antibody was omitted during the immuno-purification. **(F)** Quantification of overexpressed telomerase activity in samples from TCAB1 knock-out cells relative to parental controls (n = 4-7, mean). **(G)** Specific activity of overexpressed telomerase purified from TCAB1 knock-out cells relative to parental controls (n = 4-7, mean, T-Test). Specific activity was calculated by dividing the relative activity (see Fig. 3F) by the relative amount of TR present in immuno-purified TERT samples (Fig. 2F). The dashed lines indicate the activity level in telomerase purified from wild-type TCAB1 control cells which was normalized to 1.0.

### TCAB1 mediates telomerase assembly in living cells

The experiments presented thus far demonstrate that telomerase assembly is reduced in the absence of TCAB1 but were carried out in fixed cells or cell lysates. To analyze telomerase assembly in intact cells, we carried out live cell single-molecule imaging of 3xFLAG-HaloTag-TERT (Movie S4-5). We have previously demonstrated that there are three distinct TERT populations in the nuclei of human cancer cells (Schmidt et al., 2016): A static population (assembled telomerase RNPs bound to telomeres, Cajal bodies or other cellular structures), a slowly diffusion population, and a rapidly diffusing population (Fig. 4A). The slowly diffusing population likely includes assembled telomerase RNPs, while the rapidly diffusing particles represents TERT molecules, which are not assembled with TR (Fig. 4A). To determine the diffusion properties of TERT, when telomerase assembly cannot occur, we knocked out TR. TR knock-out was confirmed by PCR and Sanger sequencing, FISH, and qPCR (Fig. S4A-C). To define the diffusion rate of free TERT molecules, we determined the diffusion coefficient of rapidly diffusing 3xFLAG-HaloTag-TERT molecules in TR knock-out cells using the Spot-On tool, which calculates the diffusion coefficient and the fraction of particles in each TERT population by fitting the step-size distribution of TERT particles (Fig. 4A-B, Fig. S4D). Similarly, we defined the rate of diffusion of assembled telomerase RNPs by measuring the diffusion coefficient of slowly moving 3xFLAG-HaloTag-TERT molecules in control cells (Fig. 4A-B, Fig. S4D). These measurements, in TR knock-out and control cells, are the most accurate estimations possible for the diffusion coefficient of free TERT (D_fast_ = 1.9 ± 0.2 µm^2^/s) and TERT that is part of a telomerase RNP (D_slow_ = 0.35 ± 0.2 µm^2^/s), respectively. Using the diffusion coefficients, we fit the step-size distributions of TERT trajectories in control, TR knock-out, and TCAB1 knock-out cells to determine the fraction of TERT molecules in the fast, slow, and static TERT populations. Our key assumption is that rapidly diffusing TERT particles represent free TERT, while the much larger telomerase RNP is part of the slowly moving and static TERT populations. Consistent with this assumption, the fraction of TERT particles in the slowly diffusing and static populations was significantly reduced in TR knock-out cells (19 ± 4 %, 25 ± 7 %) compared to control cells (43 ± 6 %) (Fig. 4B). It is important to note that, even in TR knock-out cells, 19-25% of TERT particles are slowly diffusing or static (Fig. 4B). Because TR is absent in these cells, the slowly diffusing and static TERT molecules must be the result of interactions of TERT with cellular components other than Cajal bodies or telomeres. Importantly, we also observed a significant reduction in the fraction of slowly diffusing and static TERT particles in TCAB1 knock-out cells (29 ± 2 %) compared to control cells (43 ± 6 %) (Fig. 4B), consistent with a defect in telomerase assembly when TCAB1 is absent. The fraction of slowly diffusing and static TERT particles in TCAB1 knock-out cells (29 ± 2 %) was higher than in TR knock-out cells (19 ± 4 %, 25 ± 7 %), suggesting that telomerase assembly is strongly reduced but not completely lost in the absence of TCAB1, which is consistent with our telomerase assembly analysis using purified telomerase (Fig. 3). In addition, in the absence of TCAB1, the reduction of telomerase assembly was only detected for TERT particles that localized to the nucleoplasm but not the nucleolus (Fig. 4C-D, Fig. S4E).

**Figure 4.**
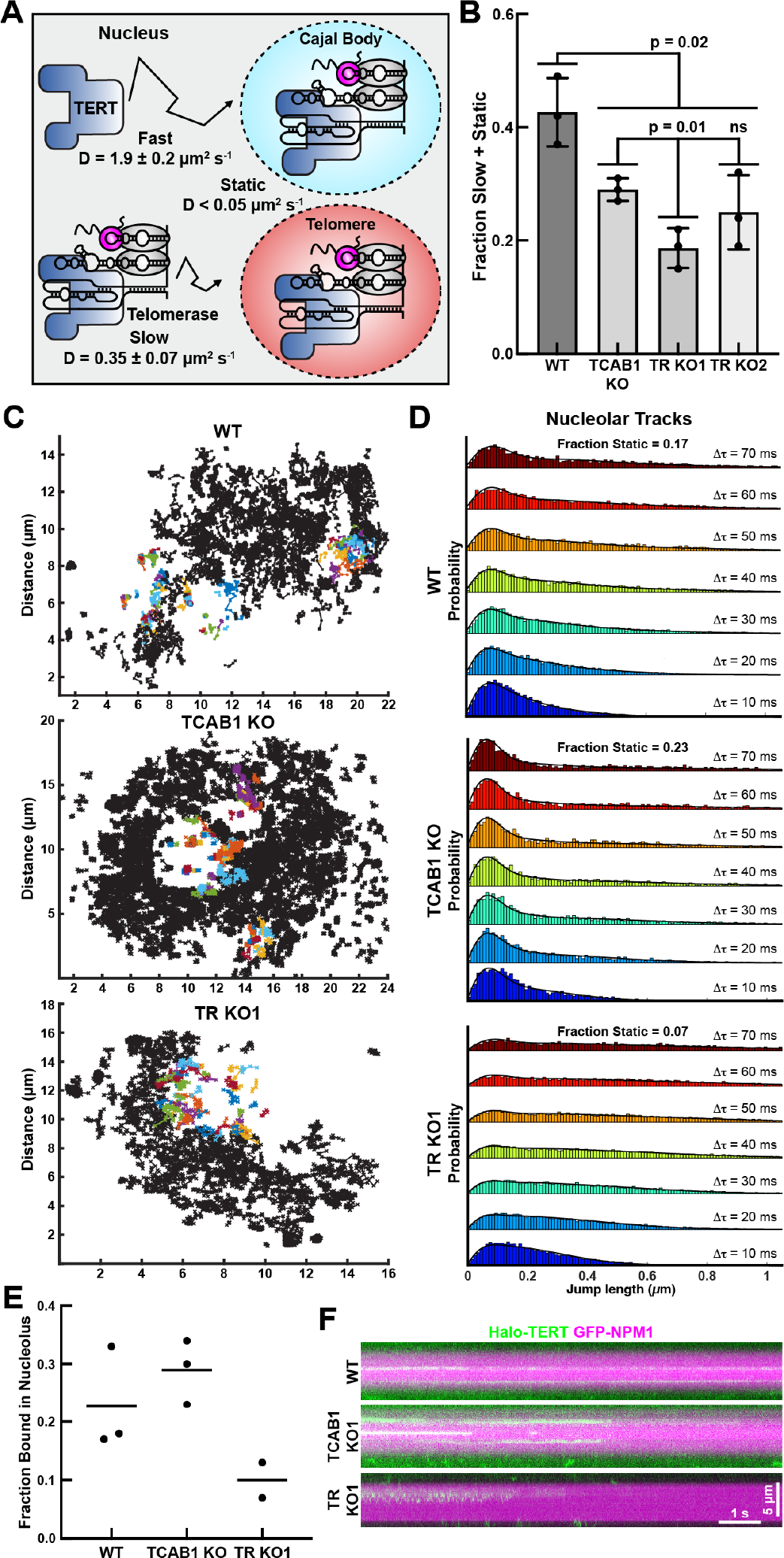
Telomerase assembly is reduced in live TCAB1 knock-out cells. **(A)** Diagram of distinct populations of TERT particles detected in control cells. Fast and slow diffusion coefficients were determined using Spot-On in control and TR knock-out cells, respectively (3 independent experiments, >15 cells per experiment per cell line, mean ± standard deviation). **(B)** Fraction of slow plus static TERT particles in control, TCAB1 knock­out, and TR knock-out cells expressing 3xFLAG-HaloTag TERT (3 independent experiments, >15 cells per experiment per cell line, mean ± standard deviation, t-test). **(C)** Graphs of HaloTag-TERT tracks that are nucleoplasmic (black) or overlap with the nucleolus for at least one frame (color) in control (WT), TCAB1 knock-out, and TR knock-out cells. **(D)** Probability density functions of the step-sizes derived from HaloTag-TERT molecules that overlap with the nucleolus for at least one frame and the corresponding 3-state model fit using the SpotOn tool (pooled data from 3 independent experiments, >18 cells total per cell line). **(E)** Quantification of the fraction of TERT particles statically bound to the nucleolus in control (WT), TCAB1 knock-out, and TR knock-out cells (2-3 independent experiments, >18 cells per experiment per cell line). **(F)** Kymographs of HaloTag-TERT particles that colocalized with nucleoli marked by GFP-NPM1 in control (WT), TCAB1 knock-out, and TR knock-out cells (also see Movies S5).

Because stable binding of telomerase to the telomere requires base-pairing of TR with the single-stranded telomeric overhang (Schmidt et al., 2018), we analyzed the interaction of TERT with telomeres. To assess TERT association with telomeres, we plotted the step size of TERT trajectories vs. the distance from the closest telomere for each step of these trajectories (Fig. S5A). In control cells, we observed an enrichment of smaller step sizes and particles in close proximity to telomeres, consistent with TERT interactions with the chromosome end (Fig. S5A). In contrast, TERT trajectories in TCAB1 knock-out cells lacked this enrichment, and the step size vs. distance from the closest telomere plots were identical to those from TR knock-out cells (Fig. S5A). Next, we filtered out TERT trajectories that came into proximity with telomeres marked by mEOS3.2-TRF2, as previously described (Schmidt et al., 2016). Diffusion analysis of these TERT trajectories using SpotOn revealed that the fraction of static TERT particles at telomeres was reduced from 12% in control cells to 4-5% in TCAB1 and TR knock-out cells (Fig. S5B). These observations indicate that, in the absence of either TCAB1 or TR, stable interactions of telomerase with telomeres occur at a lower frequency, because they require TR to stably bind to the chromosome end (Schmidt et al., 2018). Together these single-molecule imaging experiments demonstrate that in living cells, telomerase assembly is significantly reduced in the absence of TCAB1.

### TERT is not retained in the nucleolus in the absence of TR

The experiments presented thus far have demonstrated that telomerase assembly is reduced in cell lysates and in living cells when TCAB1 is absent, but do not address the sub-cellular localization of telomerase assembly. It was previously suggested that telomerase is assembled in the nucleolus and active telomerase is enriched in the nucleolus in the absence of TCAB1 (Lee et al., 2013). Alternatively, TERT could associate with TR outside of the nucleolus and a small fraction of the telomerase RNP could localize to the nucleolus after assembly. Importantly, telomerase assembly within the nucleolus would require TERT to localize to the nucleolus independently of TR. To address whether TERT can enter the nucleolus in the absence of TR, we analyzed TERT trafficking at the single-molecule level in TR knock-out cells synchronized in S-phase of the cell cycle. Nucleoli were marked by transient expression of GFP-NPM1. As described above (Fig. 1F, Movies 1-2), a very small fraction of TERT particles were detected within the nucleolus in control and TCAB1 knock-out cells. Analysis of the diffusion properties of TERT particles co-localized with GFP-NPM1 revealed that the fraction of static TERT molecules was slightly increased in TCAB1 knock-out cells (29%) compared to controls (23%, Fig. 4C-F). In contrast, static TERT molecules that co-localized with the nucleolus were less frequently detected in TR knock-out cells (10%, Fig. 4C-F, Movie S6). Interestingly, we did observe very rare TERT molecules in TR knock-out cells that were mobile but constrained within a sub region of the nucleolus which, based on our other observations, likely corresponds to TERT within the DFC that is not bound to TR (Fig. 4D, Movie S6). In total, these results demonstrate that, though rare, TERT can enter the nucleolus independently of TR but does not transition into a stably bound state that likely requires association with TR.

### TCAB1 is excluded from the nucleolus

Our experiments demonstrate that the loss of TCAB1 leads to an accumulation of TR in the nucleolus (Fig. 1). TCAB1 could lead to the depletion of TR from the nucleolus by binding to TR that has dissociated from the nucleolus and preventing it from re-entering. Alternatively, TCAB1 could bind to TR within the nucleolus and accelerate the export of TR from the nucleolus, which requires TCAB1 entry into the nucleolus. To analyze the sub-nuclear localization of TCAB1, we introduced a 3XFLAG-HaloTag at the endogenous *TCAB1* locus and generated single-cell clones that exclusively expressed 3XFLAG-HaloTag-TCAB1 (Fig. 5A). HaloTag-TCAB1 strongly accumulated at Cajal bodies (marked by BFP-coilin) and was excluded from the nucleolus (Fig. 5B, Movie S7).

**Figure 5.**
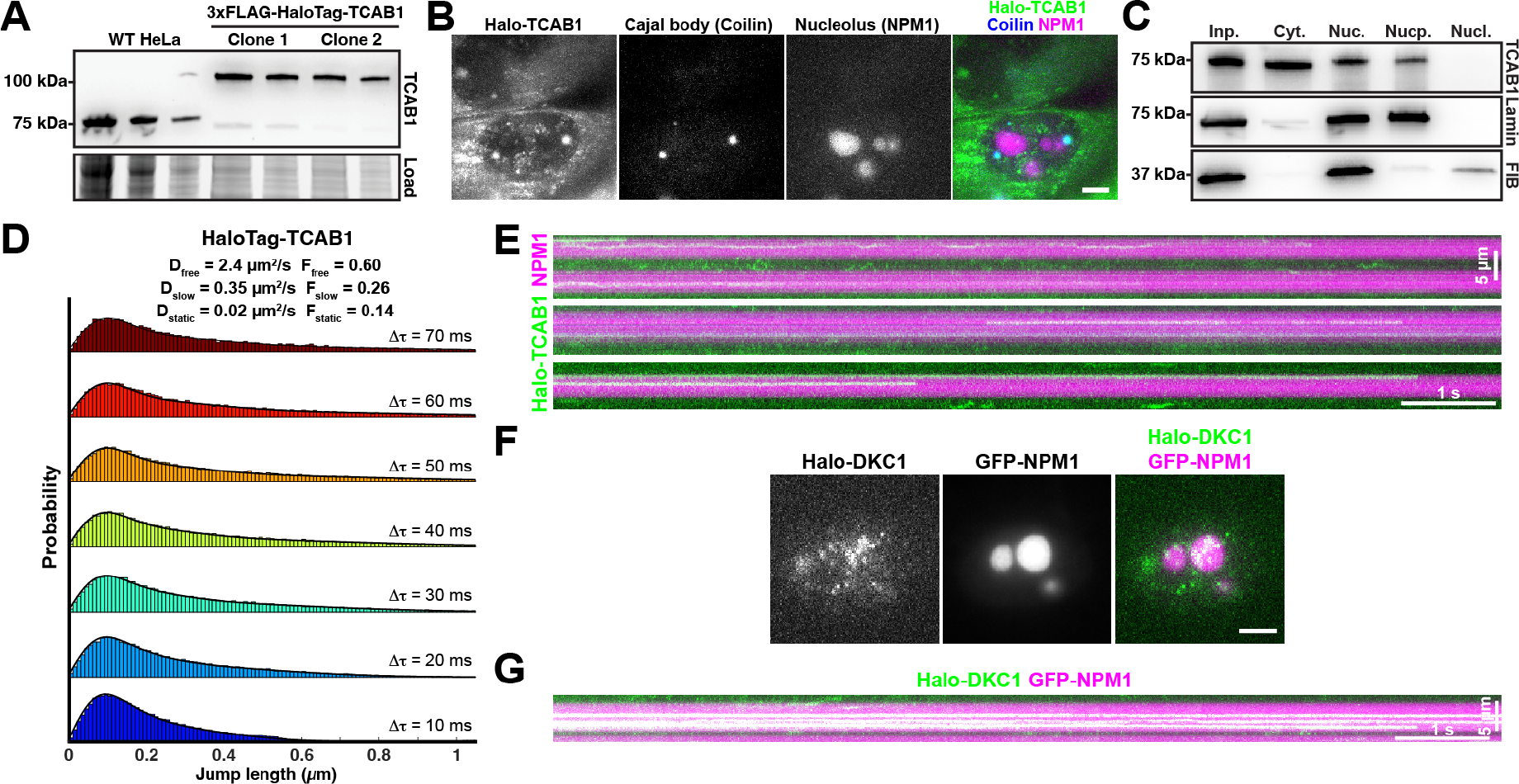
TCAB1 is excluded from the nucleolus. **(A)** Western blot probed with an antibody against TCAB1 demonstrating the exclusive expression of HaloTag-TCAB1 in genome edited Hela cells. **(B)** Maximum intensity projection of a single-molecule imaging movie of HaloTag-TCAB1 labeled with JFX650 in cells transiently expressing BFP-coilin and GFP-NPM1 to mark Cajal bodies and nucleoli, respectively (scale bar = 5 µm) **(C)** Western blots of samples of cellular fractionation experiments (left to right: Input, Cytoplasm, Nucleus, Nucleoplasm, Nucleolus) from HeLa cells. Blots were probed with antibodies against TCAB1, fibrillarin (nucleolar marker), and lamin B1 (nucleoplasmic marker). **(D)** Probability density functions of the step-sizes derived from nuclear HaloTag-TCAB1 trajectories and the corresponding 3-state model fit using the SpotOn tool (pooled data from 2 biological replicates of two independent HaloTag-TCAB1 clones, >10 cells total per replicate). **(E)** Kymograph of nucleolar HaloTag-TCAB1 particles over time. **(F)** Live cell fluorescence images of single 3xFLAG-HaloTag-dyskerin particles, nucleoli marked by GFP-NPM1 (scale bar = 5 µm). **(F)** Kymograph of nucleolar 3xFLAG-HaloTag-dyskerin particles over time.

Surprisingly, a large fraction of HaloTag-TCAB1 localized to the cytoplasm, suggesting that its import into the nucleus is restricted (Fig. 5B). To assure that the cytoplasmic localization of HaloTag-TCAB1 was not an artifact of fusing TCAB1 to the HaloTag, we carried out cell fractionation experiments, which demonstrated that untagged TCAB1 was also detected in the cytoplasm and the nucleus but was not located in the nucleolus (Fig. 5C), consistent with our imaging experiments. Single-particle tracking of HaloTag-TCAB1 within the nucleus revealed that the diffusion dynamics of TCAB1 were similar to those of TERT (Fig. 4B, Fig. 5D). Like TERT, the step size distribution of HaloTag-TCAB1 particles fit well to a three-state model (Fig. 5D). The freely diffusing state of TCAB1 had a higher diffusion coefficient than TERT (D_fast_ = 2.4 µm^2^/s vs. 1.9 µm^2^/s). Strikingly, the diffusion coefficient of the slowly diffusing state for TCAB1 and TERT were identical (D_slow_ = 0.35 µm^2^/s), consistent with TCAB1 and TERT being part of assembled H/ACA RNPs. On rare occasions, we observed HaloTag-TCAB1 molecules that associated with the nucleolus (Fig. 5E, Movie S7-8). In contrast, single HaloTag-dyskerin were readily detected stably bound to the nucleolus (Fig. 5F-G). In total, these results demonstrate that TCAB1 is largely excluded from the nucleolus and that TCAB1 molecules diffuse through the nucleoplasm at a rate similar to assembled telomerase RNPs.

### The majority of TR is tightly associated with the nucleolus in absence of TCAB1

Single-molecule imaging of TERT revealed that <10% of TERT molecules localize to the nucleolus in control and TCAB1 knock-out cells (Fig. 1G). To determine what fraction of TR associates with the nucleolus in the absence of TCAB1, we carried out cellular fractionations to isolate nucleoli. Nucleoli were purified by rupturing isolated nuclei via sonication, followed by centrifugation through a high-density sucrose cushion (Lam and Lamond, 2006). Isolated nucleoli were enriched with the nucleolar protein fibrillarin and U3 snoRNA while being depleted of lamin B1 and the 7SK RNA (Fig. 6A,B), which serve as nucleoplasmic markers, demonstrating that we successfully purified nucleoli using this approach. To determine the amount of TR found in the nucleolus and the nucleoplasm, we quantified the level of TR relative to that of the U3 or the 7SK RNA, respectively. In control cells, the majority of TR was found in the nucleoplasmic fraction, and a small amount of TR was detected in nucleoli (Fig. 6B), consistent with previous work that analyzed TR localization by live cell imaging (Laprade et al., 2020). In contrast, in TCAB1 knock-out cells, TR was depleted from the nucleoplasm and enriched in the nucleolus (Fig. 6B,C). To assess the impact of high salt concentrations, used by others to extract telomerase from human cells (Chen et al., 2018), on nucleolar integrity, we supplemented ruptured nuclei with potassium chloride, prior to isolating nucleoli. After exposure to a high salt concentration, fibrillarin and TR were found in the nucleoplasmic fraction instead of the nucleolar pellet (Fig. S6), demonstrating that nucleoli are disrupted, and TR is released when exposed to non-physiological salt concentrations. These observations confirm that >50% of TR is sequestered in the nucleolus in the absence of TCAB1 and strongly suggest that TR is tightly associated with the nucleolus under these circumstances, preventing it from entering the nucleoplasm.

**Figure 6.**
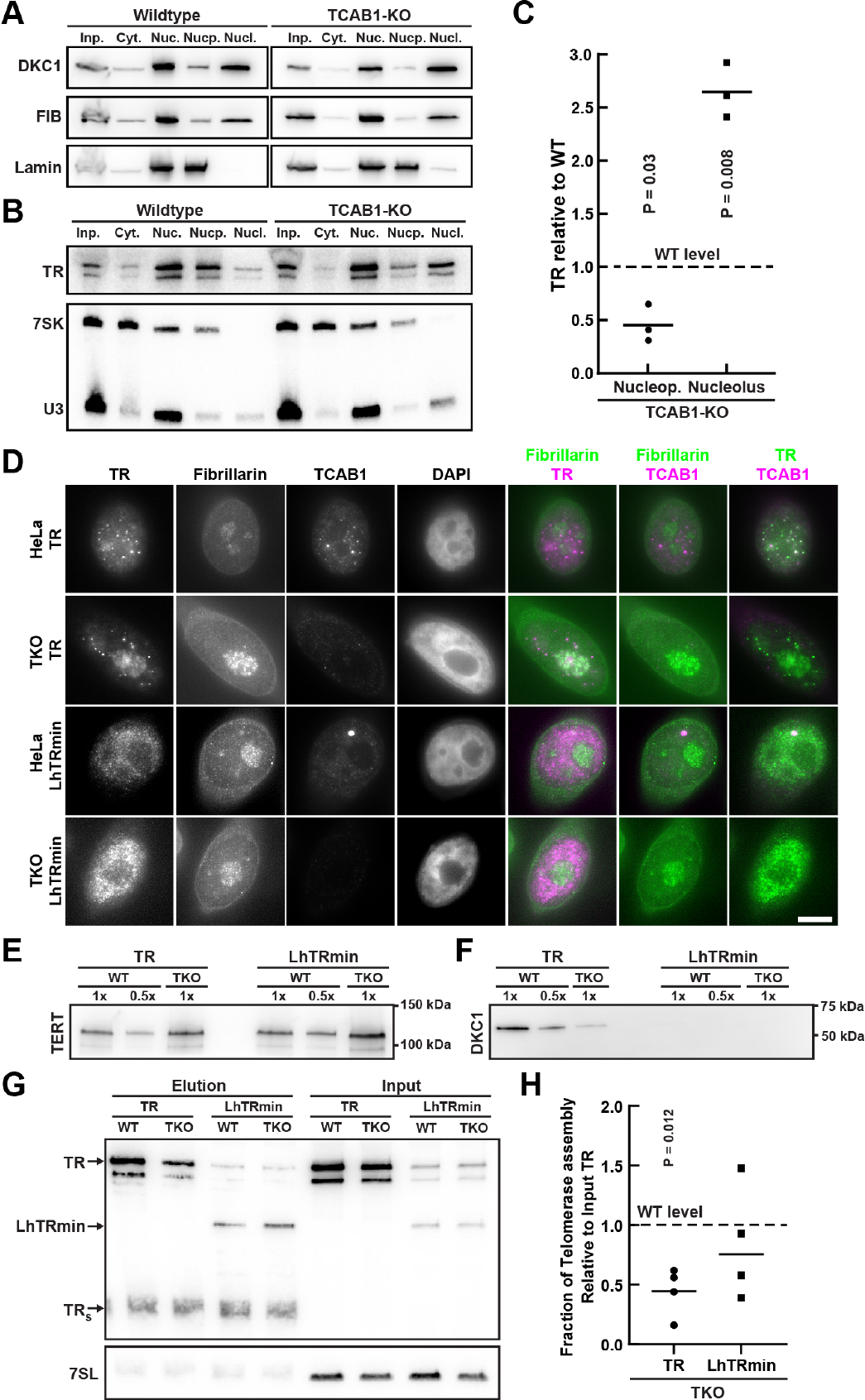
Localization of TR to the nucleoplasm rescues telomerase assembly in the absence of TCAB1. **(A)** Western blots of samples of cellular fractionation experiments (left to right: Input, Cytoplasm, Nucleus, Nucleoplasm, Nucleolus) from control and TCAB1 knock-out cells. Blots were probed with antibodies against dyskerin, fibrillarin (nucleolar marker), and lamin B1 (nucleoplasmic marker). **(B)** Northern blots of samples of cellular fractionation experiments (left to right: Input, Cytoplasm, Nucleus, Nucleoplasm, Nucleolus) from control and TCAB1 knock-out cells. Blots were first probed with radiolabeled DNA oligonucleotides complementary to TR, followed by probes complementary to the 7SK RNA (nucleoplasmic marker) and the U3 snoRNA (nucleolar marker). **(C)** Quantification of the nucleoplasmic and nucleolar abundance of TR in TCAB1 knock-out cells relative to control cells. Nucleoplasmic TR signal was normalized to the 7SK RNA signal and nucleolar TR signal was normalized to the U3 RNA signal (n = 3, mean, T-Test). **(D)** IF-FISH of control and TCAB1 knock-out cells over-expressing TERT and full-length TR or LhTRmin probed with antibodies for fibrillarin and TCAB1 and anti TR-FISH probes (scale bar = 5 µm). **(E-F)** Western blots or TERT immuno-purified from control and TCAB1 knock-out cells expressing TERT and full-length TR or LhTRmin probed with antibodies for **(E)** TERT and **(F)** dyskerin. **(G)** Northern blot of RNA extracted from input and TERT samples purified from control and TCAB1 knock-out cells expressing TERT and full-length TR or LhTRmin. Standards are *in vitro* transcribed full-length TR and truncated TR**S**. Input samples were also probed for 7SL RNA as loading control. **(H)** Quantification of TR or LhTRmin associated with TERT relative to input RNA of TERT samples purified from TCAB1 knock-out cells compared to control cells.

### Localization of the telomerase RNA to the nucleoplasm rescues telomerase assembly in the absence of TCAB1

Our data demonstrate that stable association of TERT with the nucleolus requires its interaction with TR (Fig. 4). Importantly, we cannot distinguish whether static TERT molecules within the nucleolus represent telomerase RNPs that have assembled in the nucleolus or the nucleoplasm. To address whether the telomerase RNP can assemble in the nucleoplasm, we expressed a truncate version of TR (LhTRmin) that lacks the H/ACA domain which binds to dyskerin and TCAB1 and therefore does not localize to the nucleolus or Cajal bodies (Fig. 6D) (Vogan et al., 2016). Consistent with our results shown above, assembly of TERT with full-length TR and dyskerin was significantly reduced in the absence of TCAB1 (Fig. 6E-H). In contrast, TERT association with LhTRmin was not significantly different in TCAB1 knock-out cells compared to control cells expressing LhTRmin (Fig. 6E-H). Importantly, no dyskerin was associated with TERT purified from cells expressing LhTRmin (Fig. 6F). These results demonstrate that forcing TR localization to the nucleoplasm suppresses the telomerase assembly defect observed in TCAB1 knock-out cells and indicate that the telomerase RNP can form outside of the nucleolus. Importantly, these observations suggest that the sequestration of TR in the nucleolus causes the reduction in telomerase assembly observed in the absence of TCAB1.

### Analysis of nucleolar telomerase RNA and snoRNP dynamics

The results presented thus far have demonstrated that TR accumulates in nucleoli in the absence of TCAB1 by cell fractionation and in fixed cells, but we have not addressed the dynamic localization of TR in living cells. Since TR is targeted to nucleoli by dyskerin and other H/ACA RNP components, we first used dyskerin as a surrogate for TR and H/ACA snoRNPs in general. We transiently expressed HaloTagged dyskerin in parental HeLa and TCAB1 knock-out cells and analyzed dyskerin binding to the nucleolus using fluorescence recovery after photobleaching (FRAP). We identified cells with two clearly visible nucleoli, completely photo-bleached the dyskerin signal in one of the nucleoli and quantified the recovery of the fluorescence signal (Fig. 7A, Movies S9-10). The dyskerin signal recovered rapidly (t_1/2_ = 28 s) but only ∼65% of the signal was recovered after > 4 minutes (Fig. 7B-D, Movies 9-10). This indicates that there are at least two distinct populations of dyskerin molecules in the nucleolus, a rapidly exchanging population and a static population that does not dissociate from the nucleolus over the time course of this experiment (Fig. 7D). The presence of a mobile dyskerin population was confirmed by analysis of the unbleached nucleolus, which lost fluorescence signal with similar kinetics (Fig. 7A, Fig. S7A-B). Importantly, no significant difference in dyskerin dynamics were observed in TCAB1 knock-out cells compared to parental controls (Fig. 7A-D, Fig. S7A-B), which indicates that TCAB1 is not required to extract dyskerin-containing scaRNPs from nucleoli.

**Figure 7.**
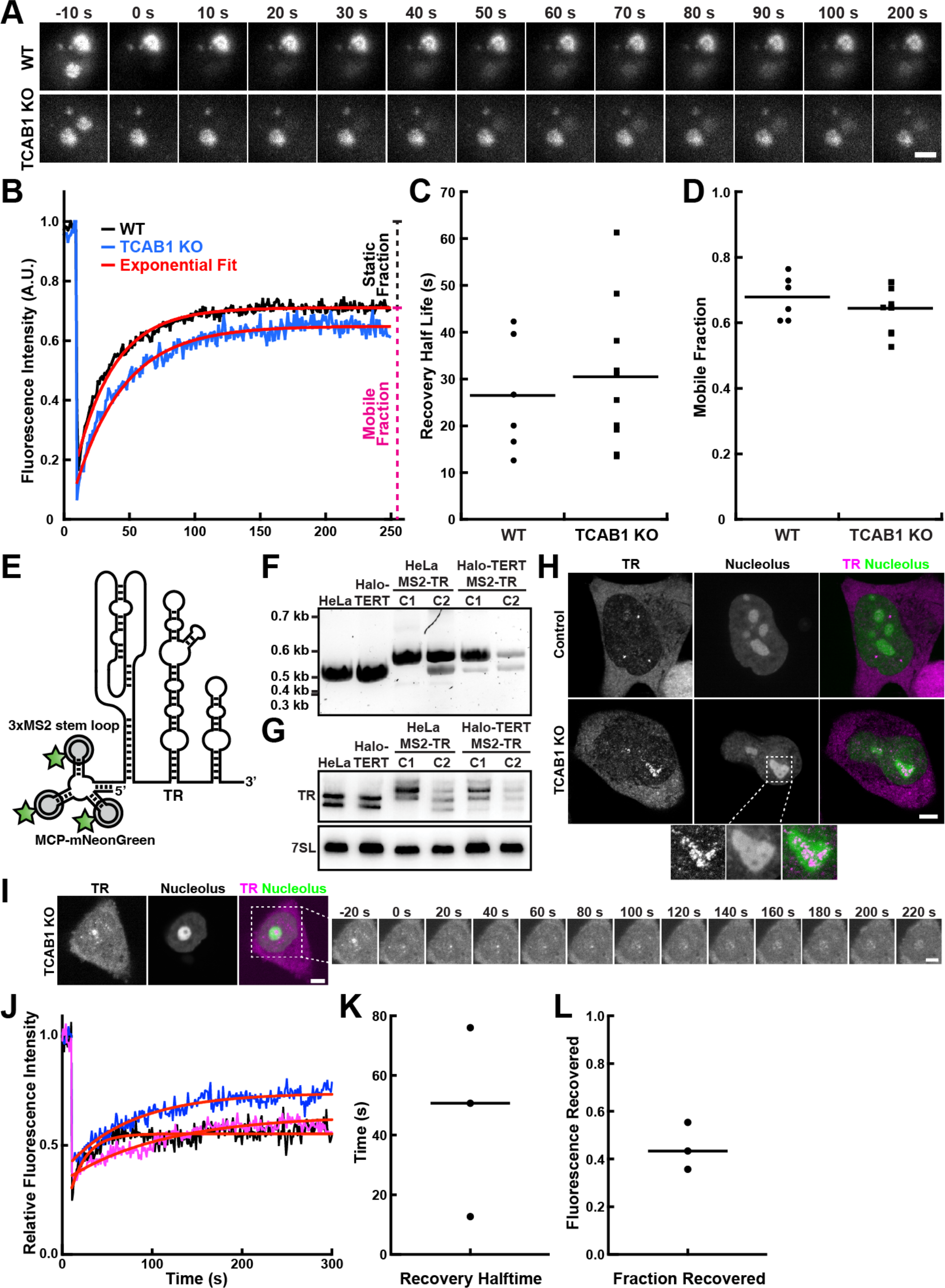
Analysis the nucleolar binding dynamics of dyskerin and TR. **(A)** Images of control and TCAB1 knock-out cells expressing 3xFLAG-HaloTag-dyskerin before and after photobleaching of nucleolar dyskerin (JFX650, scale bar = 5 µm). **(B)** Fluorescence recovery curves of nucleolar dyskerin in control and TCAB1 knock-out cells. Data was fit with a single exponential function. **(C)** Quantification of half-life of fluorescence recovery, calculated from the rate constant of the single exponential fit of the data shown in Fig. 7B (n = 6 and 9, mean). **(D)** Quantification of the mobile fraction of nucleolar dyskerin based on the single exponential fit of the data shown in Fig. 7B (n = 6 and 9, mean). **(E)** Model of 3xMS2-TR bound to MCP-mNeonGreen. **(F)** PCR analysis of the endogenous TR locus after insertion of the 3xMS2 sequence. **(G)** Northern blot probed with radioactively labeled anti-TR oligonucleolides of control cells and genome edited clones expressing 3xMS2-TR. **(H)** Live cell microscopy images of control and TCAB1 knock-out cells expressing 3xMS2-TR, MCP-mNeonGreen, and BFP-NLS to mark the nucleolus. Zoomed images demonstrate the enrichment of TR in the DFC indicated by low BFP-NLS signal within the nucleolus. **(I)** FRAP of 3xMS2-TR within the DFC of the nucleolus marked by BFP-NLS. **(J)** FRAP curves and single exponential fits of the data (red). **(K)** Quantification of the recovery halftime from single-exponential fits of the recovery curves show in (I). **(L)** Quantification of the fraction of fluorescence recovered from single-exponential fits of the FRAP curves show in (I).

To directly analyze the dynamics of TR binding to the nucleolus in the absence of TCAB1, we introduced three MS2 stem loops at the 5’ end of the endogenous TR gene (Fig. 7E-G), as previously described (Laprade et al 2020). This approach allowed us to visualize TR in living cells using the MS2 coat protein (MCP) fused to the mNeonGreen fluorescent protein, stably expressed by lentiviral transduction (Fig. 7E). Similar to endogenous TR (Fig. 1), 3xMS2-TR localized to nucleoplasmic foci in control cells (likely Cajal bodies) and to nucleoli in absence of TCAB1 (Fig. 7H, Fig. S7C). Like dyskerin (Fig. S1F), MS2-TR was specifically enriched in nucleolar regions with reduced granular component marker (BFP-NLS, Fig. S1F) intensity in the absence of TCAB1 (Fig. 7H, zoomed panel), which corresponds to the DFC of the nucleolus. To measure the dynamics of TR association with the nucleolus in TCAB1 knock-out cells, we carried out FRAP experiments of MCP-mNeonGreen bound MS2-TR clearly localized within DFC of the nucleolus (Fig. 7I). After bleaching nucleolar MS2-TR enriched in the DFC in TCAB1 knock-out cells, ∼40% of the fluorescence recovered with a halftime of recovery of ∼50 seconds (Fig. 7I-L). This demonstrates that, in the absence of TCAB1, the majority of TR is tightly associated with the nucleolus and the remaining TR molecules exchange with the nucleoplasm with kinetics comparable to dyskerin. In total, these results strongly suggest that the reduction of telomerase assembly, observed in the absence of TCAB1, is a consequence of the sequestration of TR in the nucleolus, where it infrequently encounters TERT which is largely localized to the nucleoplasm.

## Discussion

The experiments described in this study demonstrate that TCAB1 promotes telomerase assembly. In the absence of TCAB1, the telomerase RNA is targeted to the nucleolus via its association with dyskerin and other components of the H/ACA complex. In contrast to TR, TERT rarely enters the nucleolus, preventing its association with TR in cells that lack TCAB1. In addition, we demonstrate that TCAB1 does not enter nucleoli excluding the possibility that TCAB1 extracts TR containing RNPs from nucleoli. These observations suggest that nuclear compartmentalization, which is a consequence of nucleolar phase separation, counteracts telomerase assembly. Our analysis of the dynamic association of dyskerin bound snoRNPs and TR with nucleoli revealed that a significant fraction of TR containing RNPs and snoRNPs in general are tightly associated with the nucleolus. Finally, we demonstrate that while telomerase assembly is limited, the specific activity of telomerase is unchanged in the absence of TCAB1, which excludes a role of TCAB1 in telomerase catalytic function. Altogether our work supports an entirely new hypothesis for the role of TCAB1 in telomerase function in human cells and provides insight into the role phase-separated organelles play in RNA dynamics and RNP assembly and function.

### TCAB1 promotes telomerase assembly

The importance of TCAB1 for telomere synthesis is undisputed (Chen et al., 2018; Venteicher et al., 2009). Knock-out or depletion of TCAB1 results in telomere shortening (Chen et al., 2018; Venteicher et al., 2009; Vogan et al., 2016). All previous work also concluded that TR is enriched in the nucleolus in the absence of TCAB1 (Chen et al., 2018; Stern et al., 2012; Vogan et al., 2016; Zhong et al., 2011). However, prior studies conclude that TCAB1 is not required for telomerase assembly but instead plays a role in telomerase trafficking to Cajal bodies and telomeres or is required for telomerase catalysis (Chen et al., 2018; Stern et al., 2012; Venteicher et al., 2009; Vogan et al., 2016; Zhong et al., 2011). In contrast, our results demonstrate that, in the absence of TCAB1, telomerase assembly is significantly reduced. This finding is supported by two completely distinct and complementary approaches. First, we purified TERT from cell lysates and demonstrated that the fraction of TR associated with TERT is reduced to 20-40% in the absence of TCAB1 compared to control cells. Secondly, we used the single-molecule diffusion dynamics of TERT as a read-out of telomerase assembly. When telomerase assembly is impossible (in TR knock out cells), the vast majority (∼75-80%) of TERT molecules rapidly diffuse through the nucleus. In contrast, in the presence of TR, only ∼55% of TERT molecules rapidly diffuse through the nucleus, because the association of TERT with TR leads to the formation of a large RNP that moves through the nucleus at a slower rate. In the absence of TCAB1, the fraction of rapidly moving TERT molecules is significantly increased to ∼70%, consistent with a reduction in telomerase assembly. Assuming a maximal change of 25% in the fraction of rapidly diffusing TERT molecules (from 55% in wild type to 80% in TR knock-out cells), the 15% increase, observed in the absence of TCAB1 compared to wild type, corresponds to a 60% reduction (15/25) in telomerase assembly, which is consistent with our results obtained by purifying telomerase from cell lysates. In addition, we demonstrate that the TERT less frequently forms stable interactions with telomeres in the absence of TCAB1, consistent with fewer TERT molecules being bound to TR. The reduction in telomerase assembly in cells lacking TCAB1 leads to a lower number of telomerase RNPs per cell and in turn telomere shortening. Importantly, telomerase assembly is reduced but not completely abolished when TCAB1 is absent, which is sufficient to support continuous cell proliferation with a short telomere length set point.

### TCAB1 is not required for telomerase catalysis

Previous work by others has reported conflicting results regarding the role of TCAB1 in telomerase catalysis, ranging from full enzymatic activity in initial reports to substantial activity defects in the most recent study (Chen et al., 2018; Venteicher et al., 2009; Vogan et al., 2016). Importantly, both our work and the only other study that analyzed the role of TCAB1 in telomerase activity using the “gold-standard” direct telomerase extension assay concluded that telomerase activity is significantly reduced in the absence of TCAB1. While both studies concur on the degree to which telomerase activity is reduced in the absence of TCAB1, the proposed underlying molecular mechanisms differ. Chen *et al*. propose that TCAB1 is required for proper folding of the CR4/CR5 region of the telomerase RNA, which directly associates with TERT, without affecting telomerase assembly (Chen et al., 2018). Recent structural analysis of the telomerase RNP from human cells revealed that TCAB1 is located far away from the CR4/CR5 region of TR (Ghanim et al., 2021). Although it is possible that telomerase can adopt additional conformations, based on the currently available structural information it is difficult to rationalize a molecular mechanism by which TCAB1 could specifically promote CR4/CR5 folding. In addition, due to the misfolding of TR telomerase was proposed to adopt a low activity state in the absence of TCAB1. Experimentally such a low activity state would manifest itself as a reduction in the specific activity of telomerase (telomerase activity per assembled telomerase RNP). Our experiments strongly suggest that, while telomerase assembly is reduced in the absence of TCAB1, the limited amount of telomerase that can assemble is close to fully active (i.e. does not have reduced specific activity). One possible explanation for the discrepancies between the work by Chen *et al*. and our study is the methodology used to generate cell lysates. Our results demonstrate that the high salt concentration used by Chen *et al*. to generate nucleolar extracts dissolves nucleoli and releases TR (Fig. 6). Consistent with this observation, salt concentrations > 250 mM have been shown to disrupt the phase separation phenomena underlying the formation of the nucleolus (Feric et al., 2016). Solubilization of the nucleolus and release of TR would override the localization of TR and TERT to distinct sub-cellular compartments and could allow telomerase to assemble in the nuclear extract. Importantly, our single-molecule imaging of TERT dynamics is consistent with a reduction of telomerase assembly in intact cells. Altogether, our enzymatic analysis, and the positioning of TCAB1 within the telomerase RNP do not support a role of TCAB1 in TR folding and telomerase catalysis but are fully consistent with TCAB1 promoting telomerase assembly.

### Location and molecular mechanism of telomerase assembly

The sub-cellular location and order in which telomerase RNP components associate with TR in human cells have largely been unknown. It has been suggested that telomerase assembly occurs in the DFC of the nucleolus (Lee et al., 2013) . While this study clearly demonstrates that catalytically active telomerase can localize to the DFC of the nucleolus (Lee et al., 2013), it fails to show that telomerase is assembled in the nucleolus. Results by others have demonstrated that eliminating Cajal bodies does not impact telomerase activity or telomere maintenance, suggesting that Cajal bodies are not necessary for telomerase assembly (Chen et al., 2018, 2015; Vogan et al., 2016). Our single-molecule live cell imaging of TERT demonstrates that TERT is almost exclusively localized to the nucleoplasm. Since Cajal bodies are dispensable for telomerase assembly, and TERT is largely excluded from the nucleolus, we believe telomerase assembly occurs in the nucleoplasm. This model is further supported by our observation that expression of LhTRmin, which lacks the binding sites for dyskerin and TCAB1 and therefore accumulates in the nucleoplasm and is excluded from nucleoli in human cancer cells (Vogan et al., 2016), rescues the telomerase assembly defect observed in the absence of TCAB1. However, it is important to point out that we do observe a very small number of TERT molecules that localize to nucleoli in control and TCAB1 knockout cells. These static nucleolar TERT molecules require the presence of TR and therefore likely represent TERT associated with TR. Unfortunately, we are unable to determine whether this assembly of TERT and TR occurred in the nucleolus or prior to import of an assembled telomerase RNP into the nucleolus.

TCAB1 could counteract TR accumulation in the nucleolus by two possible mechanisms. Either TCAB1 could prevent the entry of TR into the nucleolus, increasing the nucleoplasmic concentration of TR and the likelihood of assembly with TERT. Alternatively, TCAB1 could extract telomerase assembled in the nucleolus by accelerating the export of telomerase from the DFC. Because TCAB1 rarely enters the nucleolus, we believe that it is unlikely that it contributes to the export of scaRNPs from the nucleolus. In addition, if the presence of TCAB1 accelerated the dynamic exchange of dyskerin-bound scaRNPs between the nucleolus and the nucleoplasm, we would expect dyskerin to bind more tightly to the nucleolus in the absence of TCAB1. Instead, our FRAP experiments demonstrate that the association of dyskerin with the nucleolus is unchanged in the absence of TCAB1.

Altogether, our observations support our model that TCAB1 promotes telomerase assembly by counteracting TR accumulation in the nucleolus to facilitate its assembly with TERT in the nucleoplasm. Our analysis of the dynamics of nucleolar TR suggests that it is tightly associated with the DFC when it is not bound by TCAB1. Since TCAB1 rarely enters the nucleolus, we believe the primary role of TCAB1 is to prevent the entry of TR into the nucleolus. In the absence of TCAB1 the equilibrium of TR localization is shifted towards the nucleolus, because re-entry is not inhibited by TCAB1 and the dissociation of TR from the DFC is very slow. The resulting nucleolar accumulation of TR effectively reduces the amount of TR available to assemble with TERT. Importantly, a very small number of TERT molecules bound to TR can be imported and trapped in the nucleolus, which would further decrease the amount of telomerase available to elongate telomeres.

### Regulation of RNP assembly by nucleolar phase-separation

In addition to the mechanistic insight into the role of TCAB1 in telomerase function, our results also demonstrate that nucleolar phase separation can effectively regulate telomerase RNP assembly in the nucleus of human cells. How RNA molecules are specifically recruited into, excluded from, or expelled from non-membrane bound organelles is a key unanswered question. One model suggests that gradual replacement of non-specific, multivalent interactions of pre-ribosomal RNAs with nucleolar proteins such as NPM1 and fibrillarin, with specific, high-affinity interactions with ribosomal proteins leads to the ejection of mature ribosomal subunits from the nucleolus (Riback et al., 2020). In this model a key driving force for the retention of RNA in the nucleolus is regions of RNA not yet bound by ribosomal proteins, that are available to interact with nucleolar proteins (Riback et al., 2020). By analogy, this model would explain why TR bound by the H/ACA complex but not associated with TERT would be sequestered in the nucleolus. Our experiments demonstrate that 60% of nucleolar TR is tightly associated with the nucleolus while the remaining 40% can exchange with the nucleoplasm, which closely corresponds to fraction of telomerase assembly observed in TCAB1 knock-out cells. We believe that TR bound to TERT can exchange with the nucleoplasm, but TR not associated with TERT remains trapped in the nucleolus. In addition to the interactions formed by the H/ACA complex with nucleolar proteins and RNA, the regions of TR that are bound by TERT in the context of telomerase (i.e. the pseudoknot, template, and CR4/CR5) would be available to form non-specific, multivalent interactions with nucleolar proteins to strengthen the association of TR with the nucleolus and prevent its release when it is not associated with TERT.

TR is a unique among the scaRNAs, because it contains the additional domains that associate with TERT. Most other box H/ACA scaRNAs are substantially shorter (<150 nucleotides) than TR (451 nucleotides), and do not contain large regions that are not bound by proteins and could form non-specific interactions with nucleolar proteins. It is, therefore, possible that in cells lacking TCAB1, TR is strongly retained in the nucleolus while other scaRNAs are less tightly bound, because they lack additional interaction sites with nucleolar proteins. Consistent with this hypothesis, we observe multiple populations of dyskerin with distinct binding dynamics in nucleoli. The weakly bound population could include dyskerin bound to scaRNAs, that are not retained in the nucleolus because their RNA targets, which would provide an additional interaction site, are not present in nucleoli. In contrast, dyskerin bound to snoRNAs would strongly associate with the nucleolus, because they also bind to their target RNAs. This provides a potential explanation for the phenotypes observed in patients with TCAB1 mutations that suffer from dyskeratosis congenita. The patients have a clear deficiency in telomerase function (Zhong et al., 2011), but no defects in splicing have been reported, which would be the consequence of complete loss of scaRNA function and their critical role in spliceosome maturation.

The mechanism by which TCAB1 binding leads to the exclusion of TR and other scaRNA from the nucleolus remains a key unanswered question. TCAB1 interacts with a very short sequence motif in TR, which is far removed from the TR regions that associate with TERT (Ghanim et al., 2021). It is, therefore, unlikely that TCAB1 binding leads to exclusion of TR from the nucleolus by reducing the number of non-specific, multivalent interactions TR can form with nucleolar proteins. As outlined above, we believe that TCAB1 prevents localization of scaRNAs to the nucleolus, rather than extracting scaRNAs that are already localized to the DFC. One potential explanation is that TCAB1 counteracts scaRNA recruitment to the nucleolus by inhibiting the nucleolar localization signals within dyskerin (Heiss et al., 1999). Dissecting the molecular mechanism by which TCAB1 leads to exclusion of TR from the nucleolus in future studies will undoubtably shed light on the fundamental principles RNP recruitment to non-membrane bound organelles and its physiological role in cell biology.

## Acknowledgements

We would like to thank members of the Schmidt lab and J. Nandakumar for discussions and critical reading of the manuscript and Luke Lavis (HHMI Janelia Research Campus) for providing HaloTag dyes. This work was supported by grants from the NIH (R00 GM120386, R01GM141354) to J.C.S.. J.C.S. was a Damon Runyon Dale F. Frey Scientist supported (in part) by the Damon Runyon Cancer Research Foundation (DFS-24-17). S.B.C. acknowledges sustained support from the Ernest & Piroska Major Foundation.

## Author contributions

B.M.K. carried out IF-FISH experiments, telomere length by Southern blot, cell cycle flow cytometry and analysis, single molecule imaging, FRAP analysis, telomerase purifications, cellular fractionations, analyzed telomerase assembly, generated mCherry- and 3xFLAG-HaloTag-dyskerin plasmids, determined their sub-cellular localization, and edited the manuscript. G.I.P. maintained cell lines, established TCAB1 and TR knock-out cell lines and carried out IF-FISH experiments. K.A.-B. assisted in establishing the TR knock-out cell line and carried out characterization of the TR knock-out cells. S.B.C. purified and characterized the anti-TERT sheep antibody. L.H. and K.Y. characterized TCAB1 knock-out clones using Southern blots. E.M.P. generated the cell lines expressing 3xMS2-TR. J.C.S. carried out all other experiments, designed the research, analyzed data, and wrote the manuscript.

## Materials and Methods

### Cell Lines and Tissue Culture

All cell lines were based on HeLa-EM2-11ht (Weidenfeld et al., 2009) and were cultured in Dulbecco’s Modified Eagle Medium including L-glutamine (Gibco) supplemented with 10% fetal bovine serum, 100 units/ml penicillin and 100 µg/ml streptomycin at 37°C with 5% CO_2_. Live cell imaging was carried out using CO_2_ independent media supplemented with 2 mM GlutaMAX (Life Technologies), 10% fetal bovine serum, 100 units/ml penicillin and 100 µg/ml streptomycin at 37°C with 5% CO_2_. For single-molecule imaging of HaloTag-TERT cell were cultured in homemade imaging dishes made by gluing 22x22 mm Nexterion coverslips (170 ± 5 µm, Schott) onto the bottom of plastic 3.5 x 1.0 cm cell culture dishes with a hole in the middle using an epoxy adhesive. Prior to chamber assembly the coverslips were washed with 1 M KOH and 100% for 30 min each in a sonicating water bath. To enrich for cells in S-phase for live cell imaging experiments, cultures we treated with complete media including 2 mM thymidine for a minimum of 16 hours. Cells were released 2 hours prior to imaging by replacing the thymidine containing media with fresh media without thymidine. Puromycin selection was carried out at a concentration of 1 µg/ml.

### Plasmid Construction and Genome Editing

All plasmids were generated by Gibson assembly (NEB) using standard protocols or by inverse PCR. All plasmids will be made available on Addgene. All Cas9 and sgRNA expression plasmids were based on pX330 (Cong et al., 2013). The homologous recombination donor for the TR knock-out was generated by assembling the genomic sequences immediately upstream and downstream (∼500 bp each, gBlocks IDT) of the TR sequence flanking a puromycin resistance cassette into HpaI linearized pFastBac. The homologous recombination donor for the insertion of the 3xMS2-tag at the endogenous was generated by cloning a left homology arm (418 bp upstream of the *TERC* gene, gBlock IDT) followed by 3xMS2-TR and 100 bp of the genomic sequence downstream of TR, followed by an SV40-promoter driven puromycin resistance marker, and finally a right homology arm (473 bp, 101-573 bp downstream of the *TERC* gene, gBlock IDT) into pFastBac. The sgRNA were designed to cut at the junction of the 3xMS2-TR insert junctions and the PAM sites were mutated to not be present in the homologous recombination donor plasmid, to assure the recombined allele was not cut by Cas9. The lentiviral plasmids for expression of MCP-NeonGreen was generated by replacing the sfGFP sequence in the pHAGE-UBC-MCP-sfGFP plasmid (a kind gift from Agnel Sfeir) with the coding sequence for mNeonGreen. The lentivirus and cell lines expressing MCP-mNeonGreen were generated as previously described (Querido et al., 2020). The homologous recombination donor for the insertion of the 3xFLAG-HaloTag into the *TCAB1* gene was generated by cloning the 3xFLAG-HaloTag sequence including a SV40-promoter driven puromycin resistance marker flanked by LoxP-sites in between two homology arms (500 bp up and downstream from the start codon, gBlocks IDT). The 3xFLAG-HaloTag-NLS plasmids was generated by adding a 3xFLAG-tag to a previously described HaloTag-NLS plasmid (a kind gift from X. Darzacq and A. Hansen) (Hansen et al., 2018). The 3xFLAG-HaloTag-dyskerin plasmid was generated by replacing TERT in our previously described 3xFLAG-HaloTag-TERT expression plasmid with the dyskerin coding sequence (Schmidt et al., 2016). The mCherry-dyskerin plasmid was generated by replacing TERT in our previously described mCherry-TERT expression plasmid with the dyskerin coding sequence (Schmidt et al., 2014). Unless otherwise stated, transfections were carried out with Lipofectamine 2000 (Invitrogen) using the manufacturer’s instructions. For FRAP analysis of dyskerin 1x10^6^ HeLa cells were transfected with 1 µg of 3xFLAG-HaloTag-dyskerin plasmid using the Lonza 4D-Nucleofector with the SE Cell Line 4D-Nucleofector X kit (Cat. V4XC-1012) and program CN-114. For single-molecule imaging of dyskerin 1 µg of a GFP-NPM1 plasmid was included in addition to the 1 µg of 3xFLAG-HaloTag-dyskerin plasmid. GFP-NPM1 WT was a gift from Xin Wang (Addgene plasmid #17578; http://n2t.net/addgene:17578; RRID:Addgene_17578) (Wang et al., 2005). For single-molecule imaging of 3xFLAG-HaloTag-NLS, 1 µg of a GFP-Nucleolin plasmid was included in addition to the 1 µg of 3xFLAG-HaloTag-NLS plasmid. The GFP-Nucleolin plasmid was a gift from Michael Kastan (Addgene plasmid #28176; http://n2t.net/addgene:28176; RRID: Addgene_28176) (Takagi et al., 2005). For FRAP analysis of nucleolar 3xMS2-TR 1 µg of mTagBFP-Nucleus-7 was transfected into HeLa cells expressing 3xMS2-TR and MCP-mNeonGreen. mTagBFP-Nucleus-7 was a gift from Michael Davidson (Addgene plasmid # 55265 ; http://n2t.net/addgene:55265 ; RRID:Addgene_55265). TCAB1 was knocked-out using a single sgRNA or two separate sgRNA and Cas9 encoding plasmids that were transfected alongside a GFP-expressing plasmid. 24 hours after transfection single-cell clones were sorted using the GFP signal. TCAB1 knock-out clones were screened by PCR and confirmed by western blot, Southern Blotting of the *TCAB1* locus and immunofluorescence imaging. TR was knocked out by transfecting two sgRNA plasmids and a homologous recombination donor plasmid. 48 hours after transfection puromycin selection was initiated and 1 week after the initiation of selection single-cell clones were generated by dilution into 96-well plates. TR knock-out was confirmed using PCR and Sanger sequencing, fluorescence *in situ* hybridization, and RT-qPCR.

### Immunofluorescence and Fluorescence *In Situ* Hybridization Imaging

Fixed cell immunofluorescence imaging and fluorescence *in situ* hybridization was carried out as previously described (Schmidt et al., 2014). Briefly, cells grown on coverslips were fixed in PBS supplemented with 4% formaldehyde. When using the HaloTag for fluorescence detection cells were incubated with 100 nM of JF646 HaloTag-ligand for 30 min prior to fixation. Unincorporated ligand was removed by 3 washes with complete media followed by placing the cells back in the incubator for 10 min to let additional dye leak out of the cells. mEOS3.2-TRF2 was detected using the intrinsic fluorescence of green form of mEOS3.2. After removing the fixation solution using 2 PBS washes, coverslips were transferred into aluminum foil covered humidity chambers with a parafilm layer and rinsed with 1 ml of PBS with 0.2% Triton X-100. Cells were than incubated in blocking buffer (PBS, 0.2% Triton X-100, 3% BSA) for 30 minutes, followed by incubation with primary antibodies diluted in blocking buffer for 1 hour. All primary antibodies were used at a concentration of 1 µg/ml. After three washes with PBS + 0.2% Triton X-100, coverslips were incubated with secondary antibodies diluted in PBS + 0.2% Triton X-100 for 1 hour. All secondary antibodies were used at a concentration of 4 µg/ml. Cells were washed three times PBS + 0.2% Triton X-100 prior to a second fixation with PBS + 4% formaldehyde. In cases where nuclear staining was used, the first of the three washing steps also included 0.1 µg/ml HOECHST. After the second fixation steps, coverslips were dehydrated in three steps with ethanol (70%, 95%, 100%), re-hydrated in 2xSSC + 50% formamide, blocked for 1 hour in hybridization buffer (100 mg/ml dextran sulfate, 0.125 mg/ml *E. coli* tRNA, 1 mg/ml nuclease free BSA, 0.5 mg/ml salmon sperm DNA, 1 mM vanadyl ribonucleoside complexes, 50% formamide, 2xSSC) at 37°C, before incubating the coverslips in hybridization buffer supplemented with three TR probes (30 ng per coverslip, /5Cy5/GCTGACATTTTTTGTTTGCTCTAGAATGAACGGTGGAAGGCGGCAGGCCGA GGCTT, /5Cy5/CTCCGTTCCTCTTCCTGCGGCCTGAAAGGCCTGAACCTCGCCCTCGCCCCC GAGAG, /5Cy5/ATGTGTGAGCCGAGTCCTGGGTGCACGTCCCACAGCTCAGGGAATCGCGCCGCGCGC) overnight at 37°C. Probe sequences were previously described (Tomlinson et al., 2006). After hybridization coverslips were washes twice for 30 minutes in 2xSSC + 50% formamide and then mounted on slides using ProLong Antifade Diamond mounting media (Life Technologies). Microscopy was carried out using a DeltaVision Elite microscope using a 60x PlanApo objective (1.42 NA) and a pco.edge sCMOS camera. We acquired 20 Z-sections spaced by 0.2 µm, followed by image deconvolution and maximum intensity projection of the sections using the DeltaVision Softworx software.

### Single-Molecule Live Cell Imaging

Live cell single-molecule imaging was carried out on an Olympus IX83 inverted microscope equipped with a 4-line cellTIRF illuminator (405 nm, 488 nm, 561 nm, 640 nm lasers), an Excelitas X-Cite TURBO LED light source, a Olympus UAPO 100x TIRF objective (1.49 NA), a CAIRN TwinCam beamsplitter, 2 Andor iXon 897 Ultra EMCCD cameras or 2 Hamamatsu Orca BT Fusion cameras, a cellFRAP with a 100 mW 405 nm laser, and a blacked-out environmental control enclosure. The microscope was operated using the Olympus cellSense software. 3xFLAG-HaloTag-TERT was labeled for 2 min in complete media supplemented with 100 nM JF646-HaloTag ligand (Grimm et al., 2015). After removing the HaloTag-ligand with three washes in complete media, cells were placed back in the incubator for 10 min to allow unincorporated dye to leak out of the cells. Cells were then transferred into CO_2_ independent media and put on the microscope which was heated to 37°C. Single-molecule imaging was carried out at 50 or 100 frames per second using highly inclined laminated optical sheet illumination (Tokunaga et al., 2008). Movies were typically 20 seconds in length (2000 frames) and were followed by a transmitted light acquisition to visualize overall cell morphology.For single-molecule imaging of 3xFLAG-HaloTag-Dyskerin, cells were labeled with 100 pM of JFX650-HaloTag Ligand (Grimm et al., 2020) for 1 min. Imaging was carried out at 100 frames per second and images of GFP-NPM1 were taken before and after single-molecule movies of dyskerin to assure the position of the nucleolus had not shifted.

### RT-qPCR

RNA samples for RT-qPCR analysis were generated by using RNeasy Mini kits (Qiagen) using ∼2 million cells as starting material. Reverse transcription was carried out using random hexamer primers and SuperScript III reverse transcriptase (Invitrogen) according to the manufacturer’s instructions. qPCR was carried out using the Maxima SYBR Green qPCR master mix (Thermo Scientific) using primers for GAPDH and TR according to the manufacturer’s instructions. All qPCR reactions were carried out in triplicates and three independent biological replicates were analyzed.

### Southern Blotting

Southern blotting was carried out using standard protocols (Southern, 2006). Briefly, genomic DNA generated by phenol-chloroform extraction after cell lysis using TE supplemented with 0.5% SDS and 0.1 mg/ml Proteinase K, was digested with BamHI (generating a 1394 bp fragment spanning exons 1-3 of the *TCAB1* locus) and separated on a 0.8% agarose gel. The DNA was then transferred on a Hybond-N+ nylon membrane using capillary transfer. The *TCAB1* locus was detected using radioactive probes (alpha-^32^P-dCTP) generated by randomly primed DNA synthesis using an 800 bp PCR product overlapping with the 1394 bp restriction fragment as a template and Klenow polymerase (NEB). Telomeric restriction fragment analysis was carried out as previously described (Nandakumar et al., 2012).

### Western Blotting

Mini-PROTEAN TGX stain-free gels (Bio-Rad) were used for SDS-PAGE. Total protein was detected using a ChemiDoc MP (Bio-Rad) after a 45 second UV activation. Western transfer was carried out using the Trans-Blot Turbo transfer system (Bio-Rad) according to the manufacturer’s instructions using the mixed molecular weight transfer setting. Immuno-blotting was carried out using standard protocols. The C-terminal TCAB1 antibody (Proteintech, 14761-1-AP) was used at a 1:2000 dilution, the N-terminal TCAB1 antibody (Novus Biologicals, NB100-68252) was used at a 1:1000 dilution, the TERT antibody (Abcam, ab32020) was used at a 1:4000 dilution, the dyskerin antibody (Santa Cruz Biotech, sc-373956) was used at a 1:200 dilution, the fibrillarin antibody was used at a 1:2000 dilution (Novus Biologicals, NB300-269), and the lamin B1 antibody was used a 1:2000 dilution. Secondary antibodies were used at a 1:5000 dilution.

### Northern Blotting

RNA was extracted from cell lysates, cellular fractions, and purified telomerase samples using the RNeasy Mini kit (Qiagen) and eluted in 30 ul of RNase free water. Purified telomerase samples were supplemented with 10 ng of a loading and recovery control prior to RNA extraction (*in vitro* transcribed TR 34-328). 15 ul of eluted RNA was mixed with 15 ul of 2x formamide loading buffer (0.1XTBE, 25 mM EDTA, 0.1% bromophenol blue, 0.1% xylene cyanol, 93% formamide) and heated to 60 °C for 5 min. Samples were separated on a 6% TBE, 7M Urea, polyacrylamide gel (Life Technologies), and transferred to a Hybond N+ membrane (Cytiva) using a wet-blotting apparatus in 1x TBE for 2 hours at 0.5 A of constant current in the cold room. After transfer, membranes were UV-crosslinked, and pre-hybridized in Church buffer for 2 hours at 50 °C. Three DNA oligos complementary to TR (GACTCGCTCCGTTCCTCTTC, GCTCTAGAATGAACGGTGGAA, CCTGAAAGGCCTGAACCTC, CGCCTACGCCCTTCTCAGT, ATGTGTGAGCCGAGTCCTG), 7SL (GCGGACACCCGATCGGCATAGC), U3 (GCCGGCTTCACGCTCAGGAGAAAACGCTACCTCTCTTCCTCGTGG), and 7SK (GTGTCTGGAGTCTTGGAAGC) were radioactively labeled using T4 PNK (NEB) and ∼10x10^6^ cpm of each probe were added to the membrane. Hybridization was carried out at 50 °C overnight. Membranes were washed three times with 2xSSC, 0.1% SDS prior to exposure to a storage phosphorescence screen (Cytiva) which was then imaged on an Amersham Typhoon IP phosphoimager (Cytiva).

### Telomerase Expression and Purification

Cell lines were transfected in 15-cm tissue culture plates at ∼90% confluency (∼25-30x10^6^ cells) using 7.5 µg of TERT plasmid, 30 µg of TR plasmid and 75 µl of Lipofectamine 2000 in 1875 ul of Opti-MEM (Cristofari and Lingner, 2006). The same amounts were used in transfecting LhTRmin (Vogan et al., 2016) or pBS U3-hTR-500 (Addgene plasmid # 28170 ; http://n2t.net/addgene:28170 ; RRID:Addgene_28170) (Fu and Collins, 2003) plasmids, which were generously gifted by Kathleen Collins . Transfected cells were split to three 15 cm dishes 24 hours after transfection. 48 hours after transfection cells were counted, harvested, and snap frozen in liquid nitrogen. Cells were lysed in 1 ml of CHAPS lysis buffer supplemented with 5 µl of RiboLock RNase inhibitor (10 mM TRIS pH 7.5, 1 mM MgCl_2_, 1 mM EGTA pH 8.0, 0.5% CHAPS, 10% glycerol) per 100x10^6^ cells and rotated at 4 °C for 30 min. Lysates were cleared in a table-top centrifuge at 21,000xg for 15 min at 4 °C. Identical cell equivalents were used for all samples. 45 µg of anti-TERT antibody was added per ml of cleared lysate and samples were rotated for 1 hour at 4 °C. Lysates were then added to 100 µl of protein G agarose and rotated for 1 hour at 4 °C. The resin was spun down at 1000xg and washed four times with 1 ml of Buffer W (20 mM HEPES pH 7.9, 300 mM KCl, 2 mM MgCl_2_, 1 mM EDTA, 1 mM DTT, 1 mM PMSF, 0.1% Triton X-100, 10% glycerol). TERT was eluted in 100 µl of Buffer W supplemented with 5 µl of 1 mM TERT peptide by rotating for 30 min at room temperature.

### Telomerase Activity Assays

Telomerase assays were carried out in 20 µl of reaction buffer (50 mM TRIS pH 8.0, 150 mM KCl, 1 mM MgCl2, 2 mM DTT, 100 nM TTAGGGTTAGGGTTAGG oligo, 10 µM dATP, 10 µM dGTP, 10 µM dTTP, 0.165 µM dGTP [*α*-32P] 3000 Ci/mmol) including 2 µl of purified telomerase for 1 hour at 30 °C. Telomerase was incubated with the substrate oligo for 15 min at room temperature, prior to initiating the reaction by addition of dNTPs. Reactions were stopped by adding 100 µl of 3.6 M of ammonium acetate supplemented with 20 µg of glycogen and 32P 5’-end labeled loading control oligos (TTAGGGTTAGGGTTAGGG, TTAGGGTTAGGGTTAG). Reaction products were precipitated using 500 µl of ice-cold ethanol and stored at -20 °C over-night. Reaction products were spun down in a table-top centrifuge at max speed for 30 min at 4 °C, washed with 500 µl of 70% ethanol, and spun down again speed for 30 min at 4 °C. The 70% ethanol was decanted, and the reaction products were dried in an Eppendorf vacuum concentrator at 45 °C. Reaction products were resuspended in 20 µl of loading buffer (0.05XTBE, 25 mM EDTA, 0.05% bromophenol blue, 0.05% xylene cyanol, 46.5% formamide) and incubated at 95 °C for 5 min. 10 µl of each sample was separated on a 12% polyacrylamide, 7 M urea sequencing gel pre-run for 45 min at 90W. Gels were dried and exposed to a storage phosphorescence screen (Cytiva) and imaged on an Amersham Typhoon IP phosphoimager (Cytiva).

### Nucleolar Isolation

Cellular fractionation was carried out using a as previously described (Lam and Lamond, 2006). All procedures were carried out on ice and centrifugations at 4 °C. Approximately 1x10^6^ million cells were harvested by trypsinization, washed with PBS, followed by incubation in a hypotonic buffer (10 mM HEPES pH 7.9, 10 mM KCl, 1.5 mM MgCl_2_, 0.5 mM DTT) to swell the cells. A small fraction of the swollen cells was collected as input sample. Swollen cells were then ruptured using pre-cooled dounce homogenizer and the tight pestle (VWR Cat. 62400-595). The ruptured cells were centrifuged at 218xg for 5 min to pellet nuclei. Nuclei where then resuspended in buffer S1 (0.25 M sucrose, 10 mM MgCl_2_), layered on top of buffer S2 (0.35 M sucrose, 0.5 mM MgCl_2_) in a 15 ml conical tube, and centrifuged at 1430xg in a swinging bucket rotor for 5 min to further purify nuclei. Nuclei were resuspended in buffer S2 and sonicated on ice for 10 seconds at 30% power (Fisherbrand Model 505, 500W). The sonicated nuclei were then layered on top of buffer S3 (0.88 M sucrose, 0.5 mM MgCl_2_) and centrifuged at 3000xg in a swinging bucket rotor for 5 min to further purify nucleoli. The nucleolar pellet was suspended in buffer S2 and centrifuged a final time at 1430xg to yield a highly purified nucleolar pellet, which was resuspended in buffer S2. Equal fractions of input, cytoplasm, nuclei, nucleoplasm, and nucleoli samples were collected and analyzed by western and northern blots. To test the impact of salt concentration on the integrity of nucleoli, nuclei ruptured by sonication were mixed 1:1 with buffer S2 containing 40 mM HEPES pH7.9 with and without 715 mM KCl, prior layering the solution on top of buffer S3.

### Cell Synchronization and Flow Cytometry

Cell lines in 15-cm tissue culture plates at a 70% confluency were blocked with 2 mM thymidine then released from thymidine after 24 hours and harvested at four time points post release: 0, 4, 8 hours. Along with each synchronization, asynchronous population was also harvested. Cells were harvested using PBS supplemented with 5 mM EDTA then fixed with ethanol, stained with propidium iodide for DNA content, and finally filtered to generate a single cell suspension. Stained samples were run through BD Accuri C6 cytometer and cell-cycle distributions were analyzed on OriginLab.

### Single-Particle Tracking

Single-particle tracking was carried out in MATLAB 2019a using a batch parallel-processing version of SLIMfast modified to allow the input of TIFF files (kindly provided by Xavier Darzacq and Anders Hansen) (Hansen et al., 2018), an implementation of the Multiple-Target-Tracing algorithm (Sergé et al., 2008), with the following settings: Exposure Time = 10 ms, NA = 1.49, Pixel Size = 0.16 µm, Emission Wavelength = 664 nm, D_max_ = 5 µm^2^/s, Number of gaps allowed = 2, Localization Error = 10^-5^, Deflation Loops = 0. Diffusion coefficients and the fraction of molecules in each distinct particle population were determined using the MATLAB version of the Spot-On tool (kindly provided by Xavier Darzacq and Anders Hansen) (Hansen et al., 2018) with the following settings: TimeGap = 10 ms or 20 ms, dZ = 0.700 µm, GapsAllowed = 2, TimePoints = 8, JumpsToConsider = 4, BinWidth = 0.01 µm, PDF-fitting, D_Free1_3State = [1 25], D_Free2_3State = [0.1 1], D_Bound_3State = [0.0001 0.1]. For all experiments we carried out 3 independent biological replicates with at least 15 cells for each cell line. The statistical significance of differences in particle fractions and diffusion coefficients were assessed using a two-tailed T-Test.

For the analysis of dyskerin trajectories a mask of the nucleolus was generated manually using the threshold function in FIJI. Dyskerin trajectories whose coordinates overlapped with the nucleolar mask for a single frame were designated as nucleolar trajectories. The remaining trajectories were designated nuclear trajectories. All data sets were analyzed using Spot-On as described above.

### Fluorescence recovery after photobleaching

Fluorescence recovery experiments (FRAP) we carried out using the same Olympus microscope used for single-molecule imaging. Cells were stained for 10 min with 100 nM JFX650-HaloTag ligand in complete media. After removing the HaloTag-ligand with three washes in complete media, cells were placed back in the incubator for 10 min to allow unincorporated dye to leak out of the cells. Cells were then transferred into CO_2_ independent media and put on the microscope which was heated to 37°C. We identified cells with two clearly visible nucleoli and bleached one of them by placing three diffraction limited bleach spots within the nucleolar 3xFLAG-HaloTag-dyskerin signal. Each spot was bleached for 100 ms at 50% laser power, which lead to complete loss of the fluorescence within the nucleolus. Cells were imaged prior to and after bleaching at 1 frame per second using the Excelitas X-cite TURBO LED light source and the 100x objective. Photobleaching due to LED exposure was negligible. To quantify FRAP we first drift corrected the movie using NanoJ (Laine et al., 2019), we then placed a region of interest (ROI) within the nucleolus and quantified mean intensity within the ROI over time.

Background signal was determined in an area of the field of view that was not covered by a cell and subtracted from the nucleolar ROI. In addition, the mean fluorescence after the bleaching pulse was divided by the fraction of total cellular fluorescence remaining after the bleaching pulse. Because the laser pulse bleaches a significant amount of total cellular fluorescence (typically 20-40%), this normalization is critical to determine the maximal amount of fluorescence recovery possible. For example, if 30% of total cellular fluorescence is lost due to the bleaching pulse, the maximal fraction of pre-bleach fluorescence than can theoretically be recovered is 70%. The recovery data was then fit using a single exponential function (1-A*e^-kt^+C), where k corresponds to the rate constant and C to the fraction of the initial signal that is not recovered (i.e. the static fraction).

For the FRAP analysis of nucleolar 3xMS2-TR bound by MCP-mNeonGreen cells were imaged on a Olympus SpinSR spinning disk confocal microscope equipped with a cellFRAP unit and a Hamamatsu ORCA Fusion BT camera. 3xMS2 TR clearly localized to the DFC of the nucleolus marked by mTagBFP-Nucleus-7 was bleached using the 405 nm laser line and recovery of the MCP-mNeonGreen was imaged every second using the 488 nm laser line. Data analysis was varied out as described above.

### Quantification of Fixed Cell Imaging

For the quantification of cellular TR distribution in control and TCAB1 knock-out cells we assigned cells into one of three categories: cells with TR only at telomeres, cells with TR only in nucleoli, and cells with TR at telomeres and in nucleoli. We carried out 3 independent biological replicates and counted a minimum of 100 cells for control and TCAB1 knock-out cells.

### Quantification of RT-qPCR data

RT-qPCR experiments were carried out in triplicate and the TR Ct value was normalized to the GAPDG Ct value. The mean *Δ*Ct (Ct of TR – Ct of GAPDH) value from three independent experiments and the corresponding standard deviation were plotted.

### Quantification of Western Blots, Northern Blots, and Telomerase Activity Assays

Gel images from western Blots, northern Blots, and telomerase activity assays were analyzed using ImageQuant TL 8.2. To quantify TR levels in Northern blots the TR band intensity was normalized to the loading and recovery control signal added to the RNA sample prior to RNA purification. To quantify telomerase activity, the whole lane intensity starting at repeat 1 was determined and divided by the sum of the loading control signals. Telomerase processivity was calculated by dividing product intensity > 7 repeats by the total signal in the respective lane. The statistical significance of the observed differences was calculated using a two-tailed T-test using a minimum of three biological replicates. Each biological replicate (independent telomerase expression and purification) was analyzed in technical triplicate.

## Supplementary Information

**Supplementary Figure 1.**
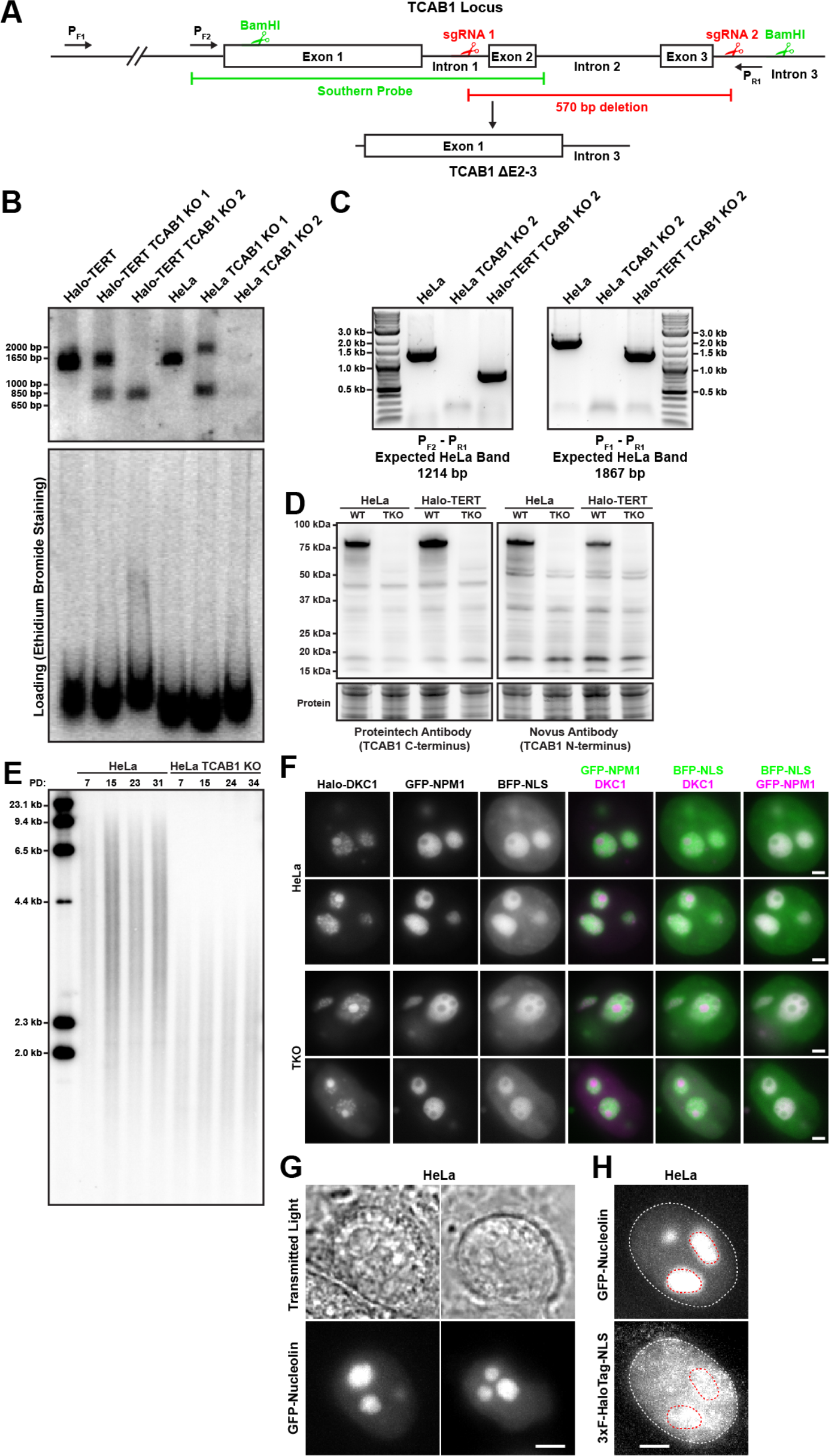
**(A)** Strategy to knock-out TCAB1 using Cas9 and two sgRNAs targeting introns 1 and 3. **(B)** Southern blot of genomic DNA digested with BamHI from parental cells and TCAB1 knock-out clones using a probes generated from a PCR product of the TCAB1 gene indicated in **(A)** demonstrating the expected truncation of the *TCAB1* gene in Halo-TERT TCAB1 KO 2. HeLa TCAB1 KO 2 carries larger deletions completely removing exons 1 and 2 from the *TCAB1* gene. **(C)** PCR using primers indicated in **(A)** of genomic DNA from parental cells and TCAB1 knock-out clones confirming the deletion of critical regions of the *TCAB1* gene show in **(B)**. **(D)** Western blots demonstrating the absence of TCAB1 protein in TCAB1 knock-out cell lines generated in HeLa and Halo-TERT cells lines using two antibodies targeting the N-terminus and C-terminus of TCAB1. **(E)** Telomere length analysis by Southern blot of telomeric restriction fragments, indicating that telomeres in TCAB1 knock-out cells are short but stable in length. **(G)** Images of HeLa cells transiently expressing GFP-nucleolin to mark nucleoli. The GFP-nucleolin signal overlaps with circular shapes visible under transmitted light illumination (scale bar = 2 µm). **(H)** Images of HeLa cells transiently expressing GFP-nucleolin and 3xFLAG-HaloTag-NLS labeled with JF646. The 3xFLAG-HaloTag-NLS signal (maximum intensity projection of 1000 frames of a single-molecule imaging movie) clearly overlaps with the GFP-nucleolin signal (red dashed outline), demonstrating that 3xFLAG-HaloTag-NLS can enter the nucleolus (scale bar = 2 µm).

**Supplementary Figure 2.**
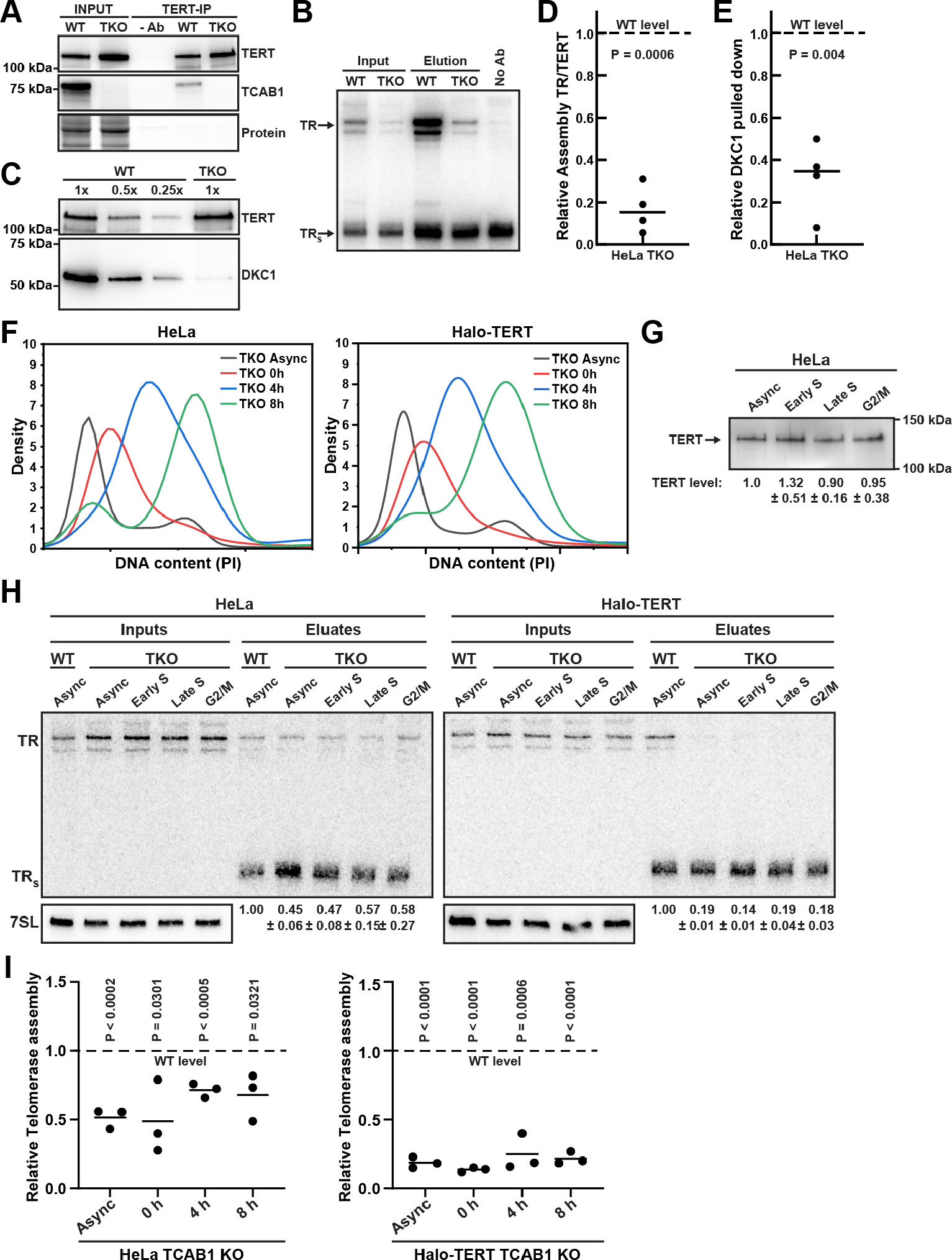
**(A)** Western blots analyzing TERT immuno-purification (using a sheep anti-TERT antibody) from Halo-TERT cells overexpressing TERT and TR probed with a rabbit anti-TERT antibody (Abcam) and a TCAB1 antibody. **(B)** Northern blot of RNA extracted from input and purified TERT samples from Halo-TERT cells overexpressing TERT and TR probed with radiolabeled DNA oligonucleotides complementary to TR. Standards are *in vitro* transcribed full-length TR and truncated TRS. TRS was added to samples prior to RNA extraction as loading and recovery control. **(C)** Western blots to analyze immuno-purified telomerase RNP composition from Halo-TERT cells. A single membrane was cut into two pieces that were probed with TERT and dyskerin antibodies, respectively. **(D-E)** Quantification of the amount of **(D)** the ratio of TR to TERT (n = 4), and **(E)** dyskerin (n = 4) in TERT purifications from Halo-TERT TCAB1 knock-out cells overexpressing TERT and TR compared to parental controls (mean, T-Test). The dashed lines indicate the level in telomerase purified from wild-type TCAB1 control cells which was normalized to 1.0. **(F)** DNA content analysis (PI staining) by flow cytometry of synchronized cell populations used for telomerase purifications. **(G)** Western blots analyzing endogenous TERT immuno-purifications (using a sheep anti-TERT antibody) from synchronized control HeLa and TCAB1 knock-out cells probed with a rabbit anti-TERT antibody (Abcam). **(H)** Northern blot of RNA extracted from input and purified TERT samples from asynchronous control HeLa cells and asynchronous and synchronized TCAB1 knock-out cells probed with radiolabeled DNA oligonucleotides complementary to TR. Standards are *in vitro* transcribed full-length TR and truncated TRS. TRS was added to samples prior to RNA extraction as loading and recovery control. Input samples were also probed for 7SL RNA as loading control. **(I)** Quantification of fraction of input TR (normalized to 7SL) co-purified with TERT or HaloTag-TERT (normalized to TRs) from TCAB1 knock-out cells relative to control cells (dashed line).

**Supplementary Figure 3.**
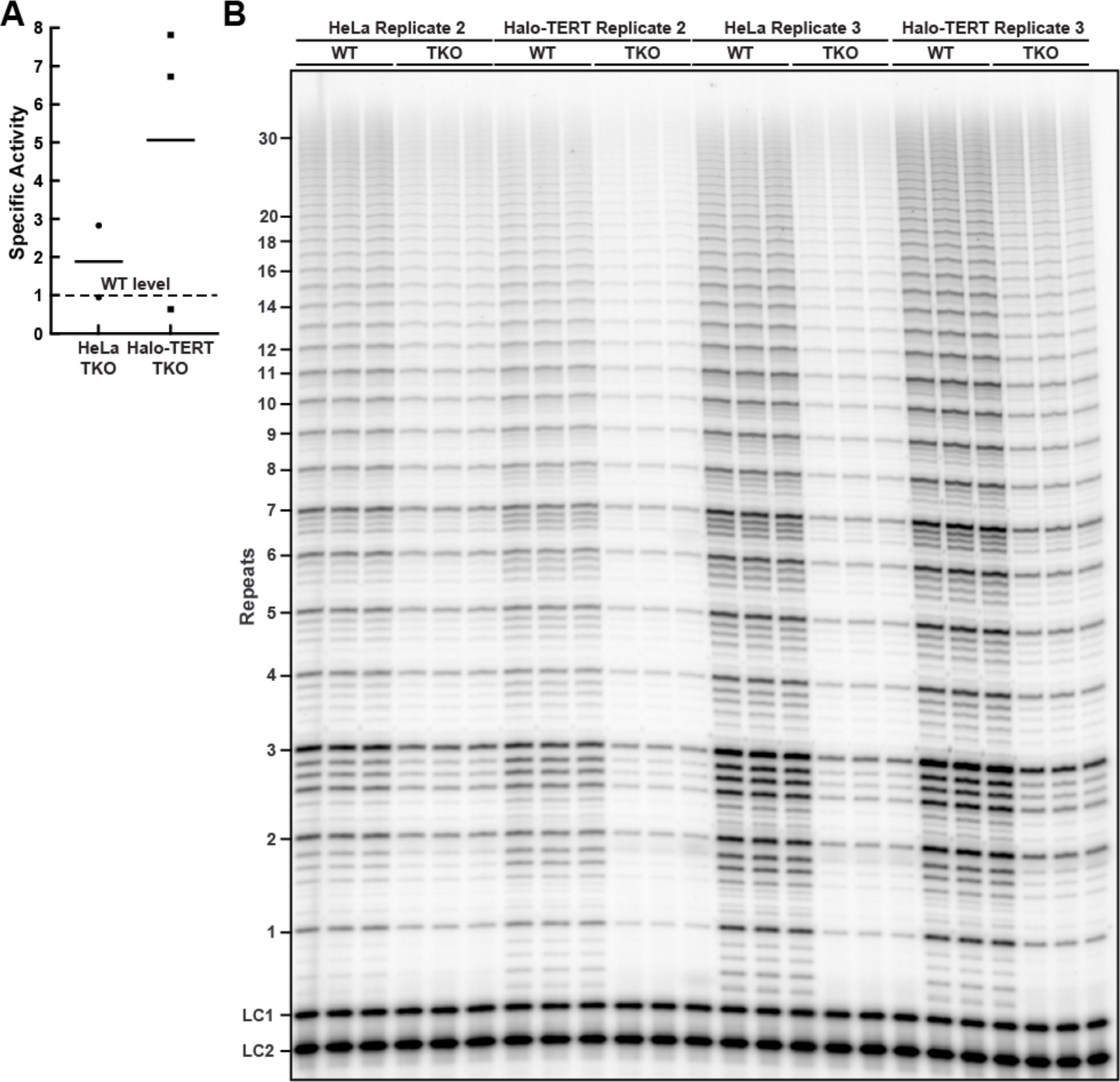
**(A)** Specific activity of endogenous telomerase purified from TCAB1 knock-out cells relative to parental controls. Specific activity was calculated by dividing the relative activity (see Fig. 5A,B) by the relative amount of TR present in immuno-purified TERT samples (Fig. 4B). The dashed lines indicate the activity level in telomerase purified from wild-type TCAB1 control cells which was normalized to 1.0. **(B)** Direct telomerase extension assay of telomerase immuno-purified from parental (WT) and TCAB1 knock-out (TKO) HeLa and Halo-TERT cell lines. LC1 and LC2, radiolabeled DNA oligonucleotide loading controls.

**Supplementary Figure 4.**
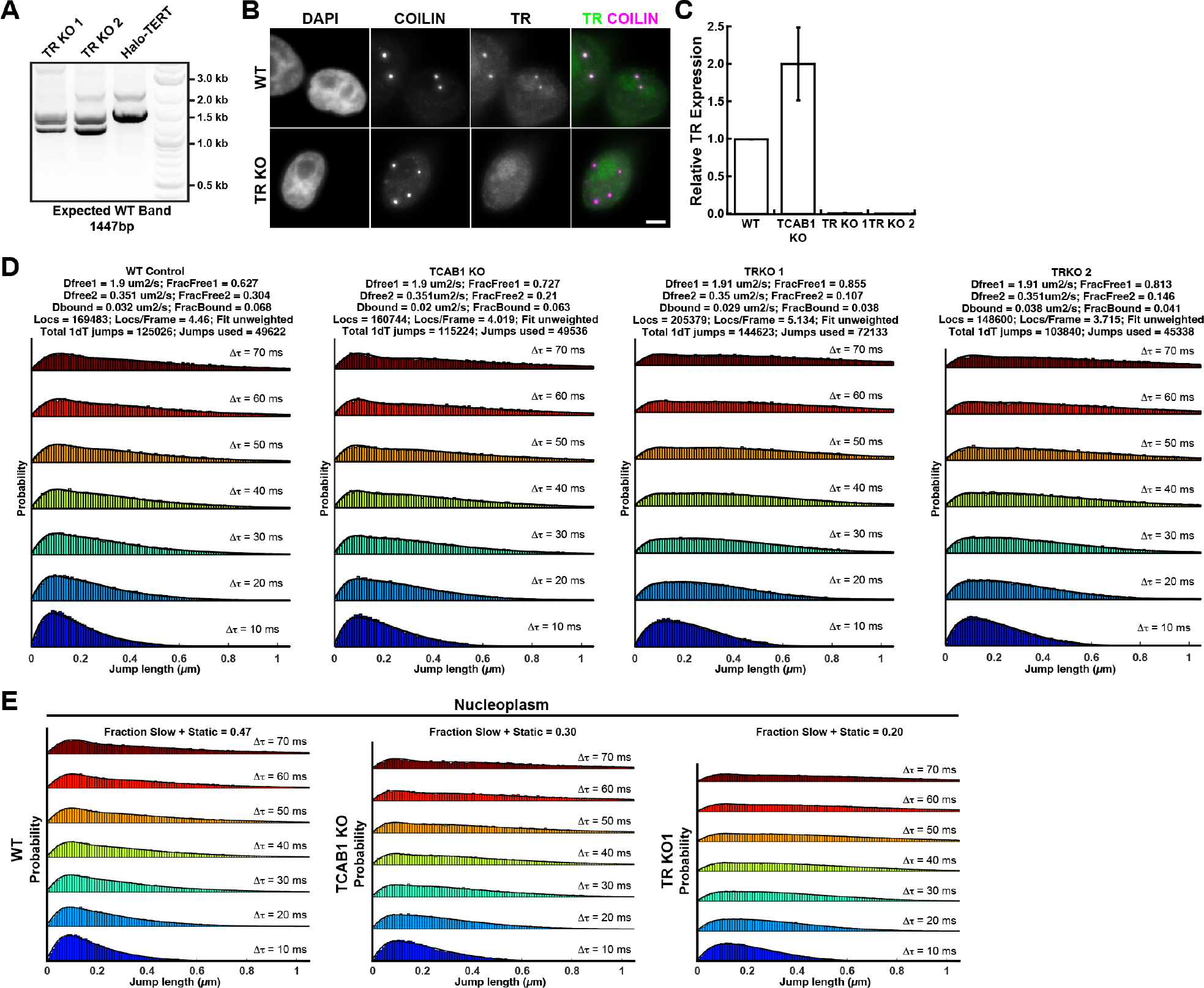
**(A)** PCR analysis of the TR locus in parental and TR knock-out clones. Both TR knock-out clones show PCR products with reduced length that were confirmed to be knock-outs by Sanger sequencing. **(B)** Images of control and TR knock-out cells probed with an antibody against coilin and FISH probes specific for TR, demonstrating the lack of TR signal in TR knock-out cells (scale bar = 5 µm). **(C)** Determination of TR levels in control, TCAB1 knock-out, and TR knock-out cells, using RT-qPCR with primers specific to TR normalized to GAPDH (3 independent biological replicates, 3 technical replicates for each biological replicate, mean ± standard deviation). **(D)** Probability density function of step sizes of HaloTag-TERT molecules from control, TCAB1 knock-out, and TR knock-out cells and the corresponding 3-state model fit using the spot on tool (Data from one of three biological replicates, >15 cells per cell line). **(E)** Probability density functions of the step-sizes derived from nucleoplasmic HaloTag-TERT molecules and the corresponding 3-state model fit using the spot on tool (pooled data from 3 independent experiments, >18 cells total per cell line).

**Supplementary Figure 5.**
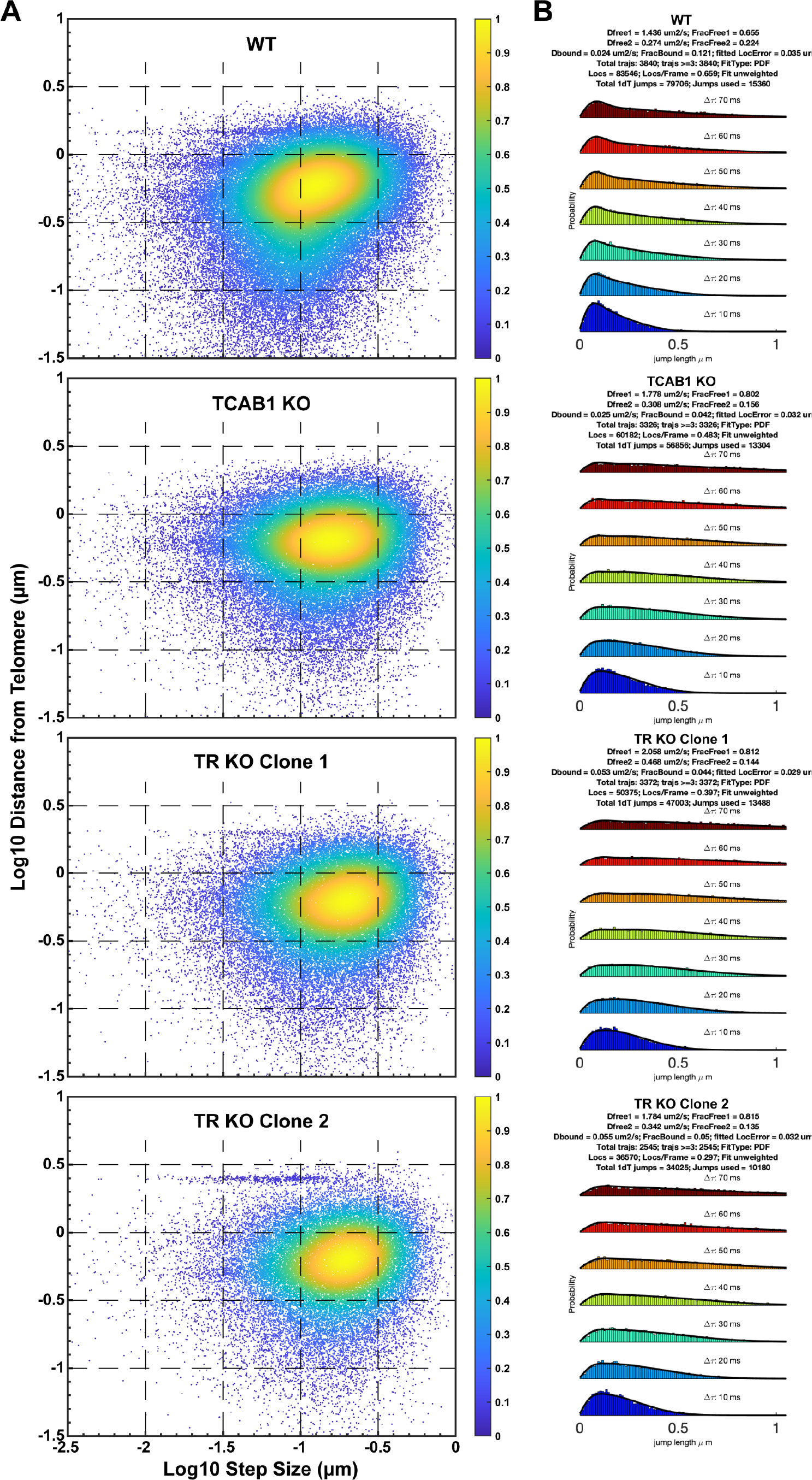
**(A)** Analysis if the step size of telomeric TERT particles relative to the distance of the particle to the closest telomere (pooled results from 3 independent biological replicates with 19-30 cells analyzed per replicate). TERT molecules bound to the telomere are expected to have small step sizes and a short distance to the closest telomere, which is apparent in the enrichment of events in the lower quadrants in the WT control. This enrichment is not observed in TCAB1 and TR knock-out cells. **(B)** Spot-On analysis of telomeric TERT particles (pooled results from 3 independent biological replicates with 19-30 cells analyzed per replicate). The fraction of bound TERT particles in TCAB1 and TR knock-out cells is 4-5%, compared to 12% in the WT control cells.

**Supplementary Figure 6.**
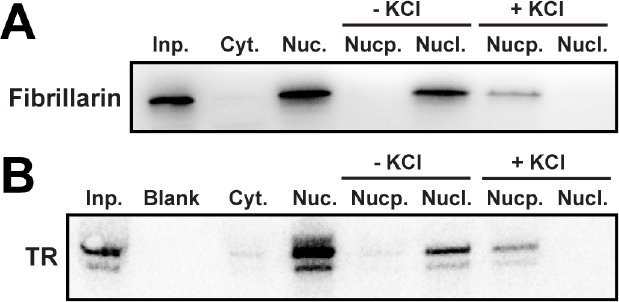
**(A)** Western blot and **(B)** Northern blot of cellular fractions from TCAB1 knock-out cells probed with an antibody against fibrillarin and probes for TR, respectively. Ruptured nuclei were either maintained at a low salt concentration or exposed to 357.5 mM KCl. The results demonstrate that nucleoli are dissolved in the presence of a high salt concentration, releasing fibrillarin and TR into the nucleoplasmic fraction.

**Supplementary Figure 7.**
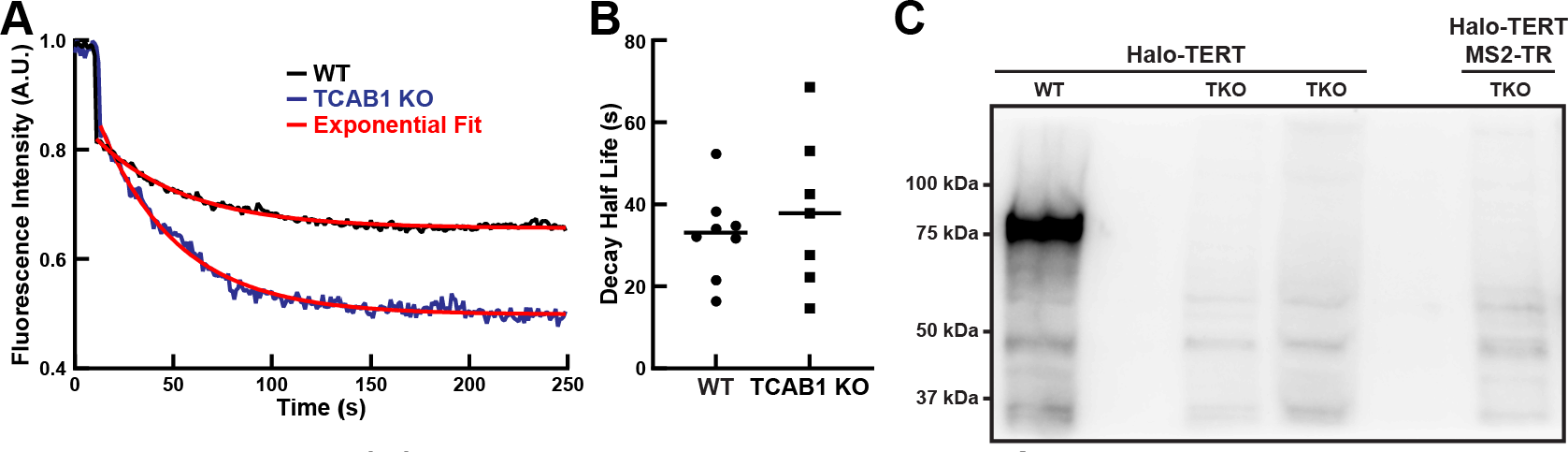
**(A)** Fluorescence recovery curves of nucleolar dyskerin in the inbleached nucleolus of control and TCAB1 knock-out cells. Data was fit with a single exponential function. **(B)** Quantification of half-life of fluorescence recovery, calculated from the rate constant of the single exponential fit of the data shown in **(A)** (n = 8 and 7, mean). **(C)** Western blot probed with an antibody against TCAB1, demonstrating the knock-out of TCAB1 in Halo-TERT MS2-TR HeLa cells.

## Movie Legends

**Movie S1.** Single-molecule imaging of 3xFLAG-HaloTag-TERT labeled with JFX650 and GFP-NPM1 to mark nucleoli in a control cell acquired at 100 frames per second second using an Andor iXon 897 Ultra camera. 200x200 pixels with a pixel size of 0.16 µm.

**Movie S2.** Single-molecule imaging of 3xFLAG-HaloTag-TERT labeled with JFX650 and GFP-NPM1 to mark nucleoli in a TCAB1 knock-out cell acquired at 100 frames per second second using an Andor iXon 897 Ultra camera. 200x200 pixels with a pixel size of 0.16 µm.

**Movie S3.** Movie of cell expressing GFP-nucleolin (red) and 3xFLAG-HaloTag-NLS (green) labeled with JF646 acquired at 100 frames per second using an Andor iXon 897 Ultra camera, showing overlap of 3xFLAG-HaloTag-NLS with nucleoli. 140x140 pixels with a pixel size of 0.16 µm.

**Movie S4.** Movie of 3xFLAG-HaloTag-TERT labeled with JF646 in a control (left), TCAB1 knock-out (middle), and TR knock-out (right) cell acquired at 100 frames per second using an Andor iXon 897 Ultra camera. Each panel is 150x150 pixels in size with a pixel size of 0.16 µm.

**Movie S5.** Single-particle tracking of 3xFLAG-HaloTag-TERT labeled with JF646 in a TR knock-out cells acquired at 100 frames per second using an Andor iXon 897 Ultra camera. Trajectories with a minimum of 5 localizations are displayed. 150x150 pixels with a pixel size of 0.16 µm.

**Movie S6.** Single-molecule imaging of 3xFLAG-HaloTag-TERT labeled with JFX650 and GFP-NPM1 to mark nucleoli in a control (left), TCAB1 knock-out (middle), and TR knock-out cells 6-hours after release from a double-thymidine block acquired at 100 frames per second using a Hamamatsu ORCA BT fusion camera. 220x220 pixels with a pixel size of 0.108 µm.

**Movie S7.** Single-molecule imaging of 3xFLAG-HaloTag-TCAB1 labeled with JFX650, GFP-NPM1 to mark nucleoli, and BFP-coilin to mark Cajal bodies acquired at 100 frames per second using an Andor iXon 897 Ultra camera. GFP-NPM1 and BFP-coilin signals were acquired before and after single-molecule imaging of 3xFLAG-HaloTag-TCAB1. 190x190 pixels with a pixel size of 0.16 µm.

**Movie S8.** Single-molecule imaging of 3xFLAG-HaloTag-TCAB1 labeled with JFX650 and GFP-NPM1 to mark nucleoli acquired at 100 frames per second using an Andor iXon 897 Ultra camera. GFP-NPM1 and BFP-coilin signals were acquired before and after single-molecule imaging of 3xFLAG-HaloTag-TCAB1. 190x190 pixels with a pixel size of 0.16 µm.

**Movie S9.** Fluorescence recovery after photobleaching of HaloTag-dyskerin labeled with JFX650 HaloTag-ligand expressed in control cells acquired at 1 frame per second.

**Movie S10.** Fluorescence recovery after photobleaching of HaloTag-dyskerin labeled with JFX650 HaloTag-ligand expressed in TCAB1 knock-out cells, acquired at 1 frame per second.

**Movie S11.** Fluorescence recovery after photobleaching of nucleolar 3xMS2-TR bound by MCP-mNeonGreen, acquired at 1 frame per second using a Hamamatsu ORCA BT fusion camera. Nucleoli were identified by transient expression of mTagBFP-Nucleus-7.

